# Oncogenic *Gata1* causes stage-specific megakaryocyte differentiation delay

**DOI:** 10.1101/791079

**Authors:** Gaëtan Juban, Nathalie Sakakini, Hedia Chagraoui, Qian Cheng, Kelly Soady, Bilyana Stoilova, Catherine Garnett, Dominic Waithe, Jess Doondeea, Batchimeg Usukhbayar, Elena Karkoulia, Maria Alexiou, John Strouboulis, Edward Morrissey, Irene Roberts, Catherine Porcher, Paresh Vyas

**Affiliations:** MRC Molecular Haematology Unit WIMM, University of Oxford, UK; Haematology Theme Oxford Biomedical Research Centre, University of Oxford, UK; Centre for Computational Biology WIMM, University of Oxford, UK; Department of Paediatrics University of Oxford, UK; Department of Hematology, Oxford University Hospitals NHS Foundation Trust, UK; Institute of Molecular Biology and Biotechnology, Foundation of Research &Technology-Hellas, Crete Greece; Biomedical Sciences Research Center “Alexander Fleming” Vari, Greece; Institut NeuroMyoGène, UCB Lyon 1, CNRS UMR 5310, INSERM U1217, Lyon, France; Cambridge Institute for Medical Research, Cambridge, UK; Department of Dentistry University of Alberta, Edmonton Alberta Canada; Rayne Institute School of Cancer & Pharmaceutical Sciences, King’s College London, UK

## Abstract

The megakaryocyte/erythroid Transient Myeloproliferative Disorder (TMD) in newborns with Down Syndrome (DS) occurs when N-terminal truncating mutations of the hemopoietic transcription factor GATA1, that produce GATA1short protein (GATA1s), are acquired early in development. Prior work has shown that murine GATA1s, by itself, causes a transient yolk sac myeloproliferative disorder. However, it is unclear where in the hemopoietic cellular hierarchy GATA1s exerts its effects to produce this myeloproliferative state. Here, through a detailed examination of hemopoiesis from murine GATA1s ES cells and GATA1s embryos we define defects in erythroid and megakaryocytic differentiation that occur relatively in hemopoiesis. GATA1s causes an arrest late in erythroid differentiation *in vivo*, and even more profoundly in ES-cell derived cultures, with a marked reduction of Ter-119 cells and reduced erythroid gene expression. In megakaryopoiesis, GATA1s causes a differentiation delay at a specific stage, with accumulation of immature, kit-expressing CD41^hi^ megakaryocytic cells. In this specific megakaryocytic compartment, there are increased numbers of GATA1s cells in S-phase of cell cycle and reduced number of apoptotic cells compared to GATA1 cells in the same cell compartment. There is also a delay in maturation of these immature GATA1s megakaryocytic lineage cells compared to GATA1 cells at the same stage of differentiation. Finally, even when GATA1s megakaryocytic cells mature, they mature aberrantly with altered megakaryocyte-specific gene expression and activity of the mature megakaryocyte enzyme, acetylcholinesterase. These studies pinpoint the hemopoietic compartment where GATA1s megakaryocyte myeloproliferation occurs, defining where molecular studies should now be focussed to understand the oncogenic action of GATA1s.

**Scientific Category:** Haematopoiesis and Stem Cells

**Key Points:** GATA1s-induced stage-specific differentiation delay increases immature megakaryocytes *in vivo* and *in vitro*, during development.

Differentiation delay is associated with increased numbers of cells in S-phase and reduced apoptosis.

## Introduction

The X-chromosome-encoded hematopoietic transcription factor GATA1 is essential for normal erythroid and megakaryocytic differentiation^1-3^. Clonal mutations acquired in fetal life, leading to loss of the N-terminal 84 amino acids of GATA1, occur in ∼28% of newborns with Down-Syndrome (DS) and are associated with either a clinically overt, or clinically silent, myeloproliferative disorder known as Transient Myeloproliferative Disorder (TMD) ^4-7^. The mutant truncated GATA1 protein is known as GATA1short or GATA1s. In most neonates with DS the mutant fetal GATA1s clone disappears by 3 months of age (^7^ and Roberts and Vyas unpublished data). In ∼3% of all neonates the TMD clone acquires additional mutations(^8,9^ that transforms the clone resulting in megakaryoblast-erythroid leukemia known as myeloid leukemia of Down Syndrome (ML-DS). Germline mutations resulting in GATA1s, in disomic individuals and families also cause disease, but rather than being oncogenic cause cytopenia^10^ including the clinical phenotype of Diamond-Blackfan Anemia^11^.

To begin to understand how GATA1s perturbs hemopoiesis, a mouse model of GATA1s has been studied^12^. These mice develop a transient megakaryoblastic myeloproliferative disorder that resolves *in utero* and likely originates from yolk sac hemopoiesis. Interestingly, these mice are anemic *in utero* leading to embryonic loss. Mice that survive then have minimal hemopoietic defects in adult life. Consistent with this human iPS cells derived from GATA1s-expressing TMD cells failed to complete erythropoiesis^13^.

This suggests that the N-terminal of GATA1 has a specific developmental role in restraining megakaryocyte production and is required for terminal red cell maturation. However, it is unclear which developmental hemopoietic cell populations require the N-terminus of GATA1 and the cellular and molecular mechanisms responsible for perturbed hemopoiesis in GATA1s cells.

To identify the cellular populations most perturbed by GATA1s, we studied hemopoietic differentiation from both ES cell culture-derived embryoid bodies (that recapitulate yolk sac hemopoiesis) and murine yolk sacs in GATA1s and control wild type GATA1 mice. We define specific stages in megakaryocyte maturation, where GATA1s megakaryocytic cells are significantly increased in overall number, exhibit decreased apoptosis, have increased numbers of cells in S-phase, exhibit a delay in terminal maturation and mature abnormally.

## Methods

### Creation of gene targeted ES cells, growth and differentiation of murine ESCs, characterisation of ES cells, flow cytometry, gene expression analysis, cell staining and microscopy, acetylcholinesterase staining quantitation, cell cycle and apoptosis assays

Details in supplemental data. Antibody clones and colors are in Supplementary Table 1. Western blotting was performed as previously described^*14*^.

### Mice

Animal studies were conducted with in accordance with the UK Home Office regulations. Embryos were processed as set out in Supplementary data.

### Statistical analyses

All experiments were performed using at least three different cultures or animals in independent experiments. The Student’s t test was used for statistical analyses. P < 0.05 was considered significant.

## Results

### Differentiation of bioGATA1 (bioG1) and bioGATA1s (bioG1s) cells

Murine bio*Gata1* and bio*Gata1*s alleles were created in male BirA ligase-expressing ES cells (ESCs)^15^ by gene targeting of X-chromosome encoded *Gata1* (**Supplemental Figure 1A-B**). Correct targeting was verified by Southern blot analysis (**Supplemental Figure 1C**) and PCR (**Supplemental Figure 1D-E**). We generated three ES cell types: BirA-bioGATA1s (hereafter, bioG1s) and as controls parental BirA (hereafter, BirA) and BirA-bioGATA1 (hereafter, bioG1).

To study the mGATA1s megakaryocyte phenotype, we used a 12 day megakaryocyte *in vitro* ES cell differentiation protocol^16^ (**Figure 1A**). ES cells were differentiated into embryoid bodies (EBs), EBs disaggregated at day 6 (d6), then CD41^+^ hemopoietic cells isolated by bead-enrichment and kit^hi^CD41^+^ cells FACS-purified (**Supplemental Figure 1F-G)** for further 6-day culture on OP9 stromal cells with cytokines to promote megakaryocyte differentiation. Western blot analysis of d6 CD41^+^ cells confirmed bioG1 cells expressed only a single higher molecular weight full-length bioGATA1 isoform, whereas bioG1s cells only expressed a single lower weight GATA1s isoform (**Figure 1B**). We next confirmed expression of *Gata1* exon 3 (common to both Gata1 and Gata1s) in BirA, bioG1 and bioG1s cells and appropriately detected cDNA spanning *Gata1* exon 2-3 only in BirA and bioG1 and not bioG1s cells (**Figure 1C**).

**Figure 1:**
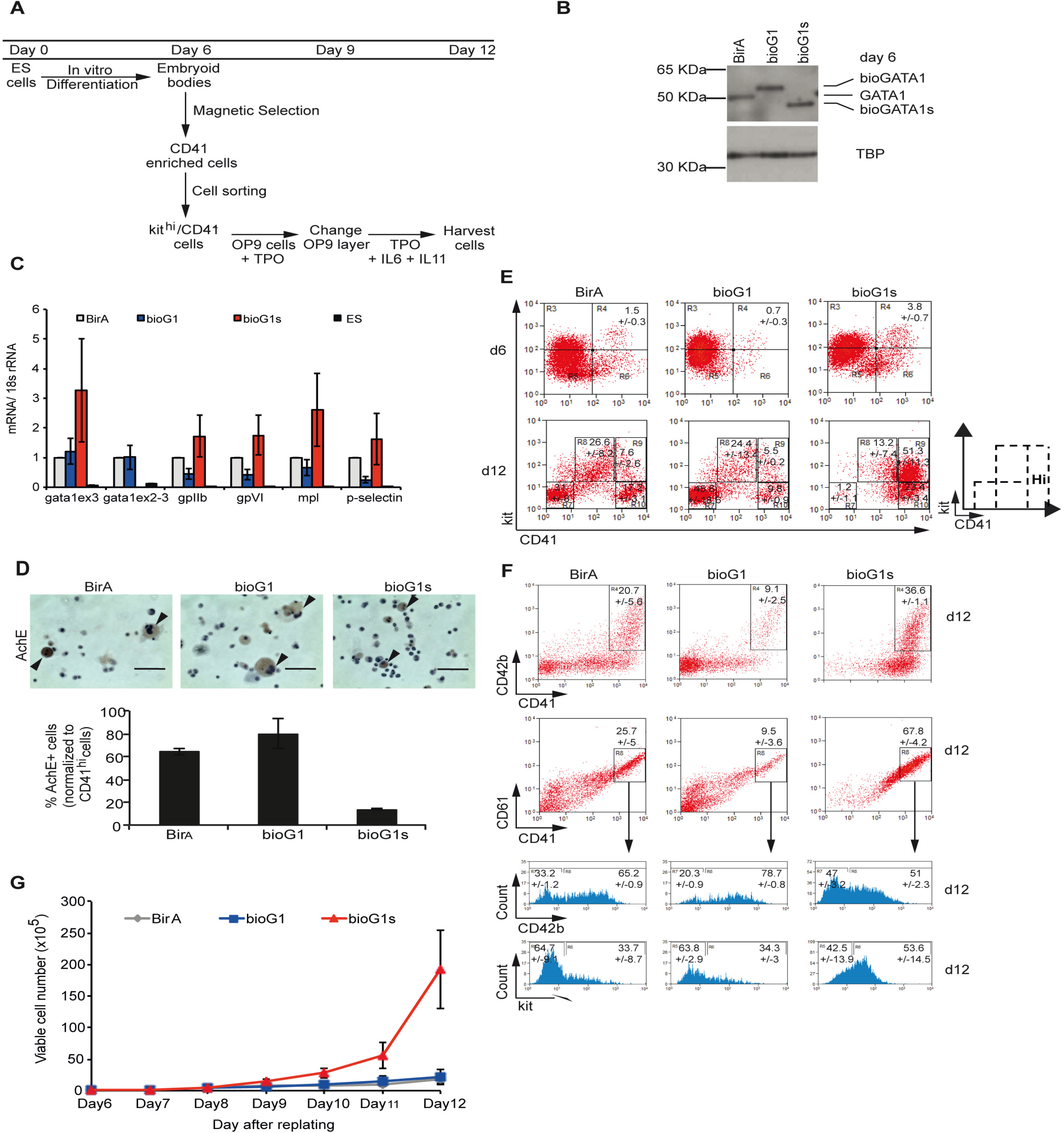
*Gata1s* ES-derived hematopoietic progenitors generate more immature megakaryocytes. A) Protocol of *in vitro* megakaryocyte differentiation from ES cells. Top, day of culture. Below, sequential steps in the culture. ES, embryonic stem cells. Tpo, thrombopoietin. IL6, interleukin 6. IL11, interleukin 11. B) Western blot probed with anti-mGATA1 antibody (top panel) and anti-TBP antibody (bottom panel) using nuclear extracts from d6 CD41^+^ cells from *in vitro* cultures. Genotype of cells is indicated above the blot. C) Expression analysis of indicated genes in 3 independent d12 EB-derived megakaryocyte cultures from BirA (grey bar), bioG1 (blue bar) and bioG1s (red bar) cells or from undifferentiated ES cells (black bar). D) Top, photomicrographs of acetylcholinesterase (AChE) stained megakaryocytes from d12 of culture (arrows). Scale bars indicate 100μm. Below, bar plot of percentages of AChE+ cells (relative to CD41^hi^ cells) in 3 different cultures. bioG1s vs BirA p=0.002; bioG1s vs bioG1 p=0.018. E) Flow cytometry showing expression of kit and CD41 on cells produced at d6 (above or d12 (below) of *in vitro* culture. Left, BirA cells, middle, bioG1 cells and right, bioG1s cells. Figures in each gate show the mean ±SD percentage of cells within the gate (5 independent experiments). Position of CD41^hi^ cells is indicated on the right of the d12 plot. d6 kit^+^CD41^+^: bioG1s vs bioG1 p=0.0002; bioG1s vs BirA p=0.005 d12 CD41^hi^: bioG1s vs bioG1 p=0.01; bioG1s vs BirA p=0.03 d12 kit^-^CD41^-^: bioG1s vs bioG1 p=0.001; bioG1s vs BirA p=0.003 F) Flow cytometry showing expression of CD42b and CD41 (top) and CD61 and CD41 (middle) at d12 of culture. Bottom, CD42b and kit expression in CD41^+^CD61^+^ cells. Left, BirA cells, middle, bioG1 cells and right, bioG1s cells. Figures in each gate show the mean ±SD percentage of cells within the gate (3 independent experiments). CD41^+^CD61^+^: bioG1s vs BirA p=0.01; bioG1s vs bioG1 p=0.004 CD42b: bioG1s vs BirA p=0.04; bioG1s vs bioG1 p=0.02 G) Viable cell count (y-axis) from d6 to d12 in culture (x-axis) when CD41^+^ cells from BirA (grey line), bioG1 (blue line) and bioG1s (red line) EBs were replated on OP9 layer with cytokines. Dead cells were excluded by trypan blue staining. d12: bioG1s vs BirA p=0.008; bioG1s vs bioG1 p=0.009

Next, we tested the lineage characteristics of cells produced by the 12 day culture. First, we took all cells from day 12 (d12) and confirmed expression of megakaryocyte genes *gpIIB, gpVI, mpl* and *p-selectin* in BirA, bioG1 and bioG1s cells but not in ES cells (**Figure 1C**). Next, by staining d12 cells with megakaryocyte-specific acetylcholinesterase stain (**Figure 1D**) we confirmed megakaryocyte production. Interestingly, bioG1s cultures produced significantly fewer megakaryocytes (bioG1s vs BirA p=0.002; bioG1s vs bioG1 p=0.018) providing a first clue that megakaryocyte differentiation is impaired by GATA1s.

To obtain a more complete initial view of megakaryocyte differentiation we analyzed kit (marker of immature hemopoietic cells) and CD41 (marker of megakaryocyte maturation) expression at d6 and d12 of culture (**Figure 1E and Supplemental Figure 1H**). D6 bioG1s EBs produced significantly more kit^+^CD41^+^ (hemopoietic) cells (bioG1s: 3.8±0.7%; BirA: 1.5±0.3%; bioG1: 0.7±0.3%) (bioG1s vs bioG1 p=0.0002; bioG1s vs BirA p=0.005). By d12, there were significantly more CD41^hi^ cells in bioG1s cultures than bioG1 and BirA cultures (bioG1s: 74.7±14.2%; BirA: 24.8±0.7%; bioG1: 15.3±0.7%) (bioG1s vs bioG1 p=0.01; bioG1s vs BirA p=0.03) but most bioG1s CD41^hi^ cells still expressed the immaturity marker, kit. Finally, there were significantly fewer non-megakaryocyte kit^-^CD41^-^ cells in bioG1s compared to bioG1 and BirA cultures (bioG1s: 1.2±1.1%; BirA: 31±8%; bioG1: 45.6±18.6%) (bioG1s vs bioG1 p=0.001; bioG1s vs BirA p=0.003).

To further characterize megakaryocyte marker expression, we confirmed that CD41^+^ megakaryocytes also co-expressed the mature megakaryocyte markers CD42b and CD61 at d12 (**Figure 1F and Supplemental Figure 1H**), paradoxically even in kit expressing cells in bioG1s cells. Interestingly, there were significantly greater percentage of CD41^+^CD61^+^ cells in bioG1s cultures compared to control bioG1 and BirA cultures (bioG1s vs BirA p=0.01; bioG1s vs bioG1 p=0.004). Finally, bioG1s cells also expressed lower levels of the maturity marker CD42b on CD41^+^CD61^+^ cells (bioG1s vs BirA p=0.04; bioG1s vs bioG1 p=0.02).

Finally, we measured cell growth by counting viable cell numbers daily from d6 to d12 (**Figure 1G**). Numbers of cells in BirA, bioG1 and bioG1s were similar from d6 to d9 but then increased significantly in bioG1s cultures and were 10-fold greater at d12 compared to both BirA (p=0.008) and bioG1 (p=0.009) cultures.

In summary, the cultures produced both megakaryocyte and non-megakaryocyte cells. Compared to wild type GATA1 hemopoietic cells, bioG1s cells were more proliferative, producing more immature megakaryocytes and fewer non-megakaryocytic cells.

To characterise the kinetics of abnormal differentiation we sampled cultures daily from d6 to d12 (**Figure 2A-D**). Starting with FACS-purified d6 kit^hi^CD41^lo^ cell population (termed P1), we monitored maturation (lower the level of kit expression the more mature the cells) and acquisition of the megakaryocyte lineage (increasing CD41 expression).

**Figure 2:**
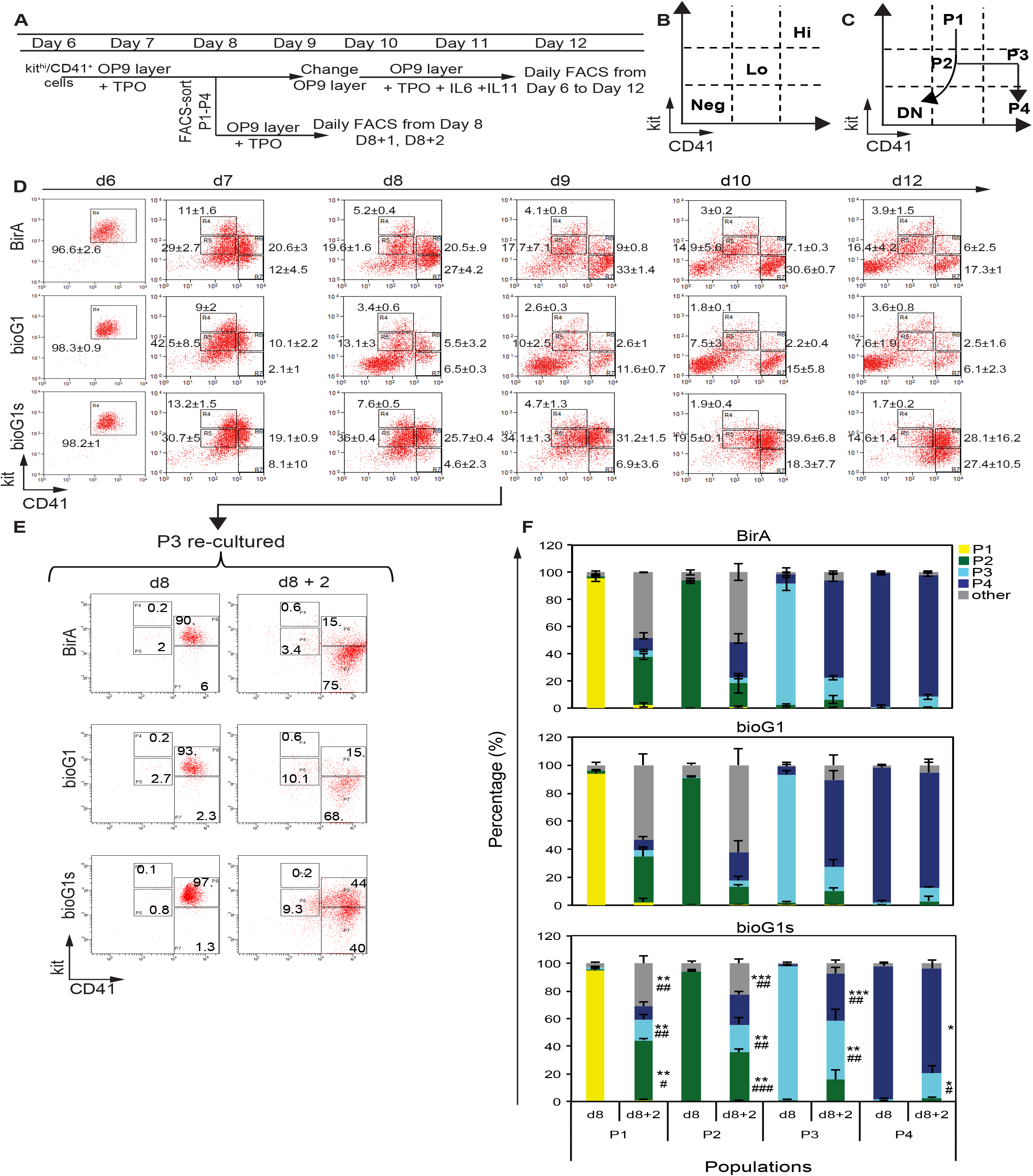
Gata1s hemopoietic cells have abnormal differentiation kinetics. A) Schematic of experiment. Hemopoietic cells (kit^hi^CD41^lo^) from BirA, bioG1 and bioG1s d6 EBs were cultured for another 6 days (up to d12). Aliquots of culture were analyzed daily for kit and CD41 expression by FACS. In parallel, at d8, populations P1-P4 (see panel B-C) cells were purified by FACS-sorting, from cultures of three genotypes, re-cultured for 2 days and kit and CD41 analyzed by FACS analysis. B) Schematic of levels of kit and CD41 expression detected by flow cytometry. Neg, negative; lo, low and hi, high. Different levels of kit and CD41 expression define different hemopoietic cell populations in panels C-E. C) Schematic summary of the data from Panel (D), showing the two branches of hemopoietic differentiation undertaken by the initial kit^hi^CD41^lo^ (P1 population). P1 cells differentiate into P2 cells (kit^lo^CD41^lo^). P2 cells then differentiate into either DN (double negative, kit^-^CD41^-^) cells or into P3 (kit^lo^CD41^hi^) cells. P3 cells differentiate into P4 (kit^-^CD41^hi^) cells. D) Representative FACS plots showing the differentiation of d6 hemopoietic cells (kit^hi^CD41^lo^, termed P1 population) from BirA (top), bioG1 (middle) and bioG1s (bottom) cultures from d7 to d12 monitored by kit and CD41 expression. Numbers within gates are the mean%±1SD of cells within the gate from 3 independent experiments. DN = day12: bioG1s vs bioG1 p=0.004; bioG1s vs BirA p=0.00009; P3 = day10: bioG1s vs bioG1 p=0.016; bioG1s vs BirA p=0.021; P4 = day9: bioG1s vs bioG1 p=0.09; bioG1s vs BirA p=0.009; P4 = day12: bioG1s vs bioG1 p=0.026; bioG1s vs BirA p=0.0015 E) Example of the re-culturing of FACS-purified d8 populations for two additional days. Here, P3 cells were FACS-purified from BirA cultures (top), bioG1 (middle) and bioG1s (bottom) cultures and re-cultured for 2 days. Left, FACS plots of post-sort purity checks of sorted P1-P4 cell populations. Right, expression of kit and CD41 expression of the 4 sorted populations after two days of culture. F) Quantitation of the different populations generated by FACS-sorted d8 P1, P2, P3 and P4 populations after an additional 2 days of culture. *p<0.05, **p<0.01 and ***p<0.001 *versus* BirA. ^#^p<0.05, ^##^p<0.01 and ^###^p<0.001 versus BirA. P1 replating = P2: bioG1s vs bioG1 p=0.024; bioG1s vs BirA p=0.002; P3: bioG1s vs bioG1 p=0.006; bioG1s vs BirA p=0.005 P2 replating = P2: bioG1s vs bioG1 p=0.00003; bioG1s vs BirA p=0.006; P3: bioG1s vs bioG1 p=0.005; bioG1s vs BirA p=0.003 P3 replating = P3: bioG1s vs bioG1 p=0.005; bioG1s vs BirA p=0.003; P4: bioG1s vs bioG1 p=0.001; bioG1s vs BirA p=0.0001

The temporal sequence of flow cytometric plots suggested that control cells (BirA and bioG1) first showed a decrease in kit expression level, generating a kit^lo^CD41^lo^ population (termed P2) (seen at d7). Cells in P2 then divided into two differentiation branches (**Figure 2B-D and Supplemental Figure 2**). In one branch, cells progressively lost expression of both kit and CD41 (d8 onwards) to generate a kit^-^ CD41^-^ population (double negative, DN, cells). This DN population was mainly composed of erythroid cells (see below). In the other branch, P2 cells also differentiated towards the megakaryocytic lineage with an increase in CD41 expression level (kit^lo^CD41^hi^ population, called P3) (d7 onwards) followed by loss of kit expression (kit^-^CD41^hi^ population, called P4) (d8 onwards).

In contrast, there were two marked differences in bioG1s cultures (**Figure 2D**). First, they generated far fewer DN cells (day12: bioG1s, 2.2±1.6%; BirA, 31.4±2.7%; bioG1, 53.3±15%) (bioG1s vs bioG1 p=0.004; bioG1s vs BirA p=0.00009). Second, bioG1s cells showed enhanced differentiation into the P3 population (d9-10) (d10: bioG1s, 39.6±6.8%; BirA, 7.1±0.3%; bioG1, 2.2±0.4%) (bioG1s vs bioG1 p=0.016; bioG1s vs BirA p=0.021) but with a delay of differentiation into P4 (best seen at d8-9, more cells in P4 in control cultures; d9: BirA, 33±1.4%; bioG1, 11.6±0.7% bioG1s, 6.9±3.6%) (bioG1s vs bioG1 p=0.09; bioG1s vs BirA p=0.009). In contrast, there were more cells in bioG1s in P4 at d12 (bioG1s, 27.4±10.5%; BirA, 17.3±1%; bioG1, 6.1±2.3%) (bioG1s vs bioG1 p=0.026; bioG1s vs BirA p=0.0015).

Though this temporal analysis was suggestive of two differentiation branches and hierarchical relationships between P2, P3, P4 and P2 and DN (**Figure 2C**) to provide more definitive proof we FACS-sorted each population (P1, P2, P3, P4) individually at d8, re-cultured them for two days. During re-culture we analyzed kit and CD41 expression in the progeny produced (**Figure 2E**, re-culture of P3, **Supplemental Figure S3A-B**, re-culture of all the populations, **Figure 2F**, data summary,). FACS-sorted P1 generated all the other populations. Purified P2 generated all the populations except P1. P3 differentiated primarily into P3 and P4 only but not DN cells. Finally, re-culturing of P4 cells generated principally P4 cells. These data were consistent with the differentiation branches and hierarchical relationships in **Figure 2C**.

Comparing the differentiation potential of FACS-sorted bioG1s populations to control BirA and bioG1 cells (**Figure 2F**), bioG1s P1 cells also generated significantly more P2 and P3 cells than BirA and bioG1 P1 cells (bioG1s P2, 43.58±1.24%; BirA P2, 36±2.3%; bioG1 P2, 32.5±7%; bioG1s P3, 14.93±3.73%; BirA P3, 4.27±0.7%; bioG1 P3, 4.47±1.05%). BioG1s P2 cells also generated significantly more P2 and P3 cells than BirA and bioG1 P2 cells (bioG1s P2, 35.43±2.45%; BirA P2, 17.83±7.25%; bioG1 P2, 13.03±1.29%; bioG1s P3, 20.03±4.96%; BirA P3, 4.23 ±1.22%; bioG1 P3, 4.43±2.76%). Finally, bioG1s P3 cells generated more P3 but fewer P4 cells compared to BirA and bioG1 P3 cells (bioG1s P3, 42.83±8.04%; BirA P3, 16±1.57%; bioG1 P3, 17.33±4.67%; bioG1s P4, 34.18±4.47%; BirA P4, 72.13 ±4.37%; bioG1 P4, 61.77±6.71%).

Taken together, kit^hi^CD41^lo^ hemopoietic progenitors differentiate either into non-megakaryocytic DN cells or megakaryocytic cells with increased CD41 expression and loss of kit expression. BioG1s mutant cells have a reduced ability to differentiate into DN cells but generate more megakaryocytic cells but with a partial differentiation delay at the P3 population stage.

### Reduced erythroid differentiation by bioGATA1s hemopoietic cells

To confirm the identity of kit^-^CD41^-^ cells (DN), we analyzed morphology, cell surface markers and mRNA expression profile of FACS-purified cells (**Figure 3A-C and Supplemental Figure 4 A-F**). Morphologically, DN cells were primarily erythroid cells, at different stages of maturation, with hardly any granulated myeloid cells (**Figure 3A**). Approximately 50% of BirA and bioG1 DN cells were Ter119^+^ and 5% were Mac1^+^, Gr1^+^ or both Mac1^+^Gr1^+^ (**Figure 3B, Supplemental Figure 4A-C**). The DN population was markedly reduced in bioG1s cultures compared to bioG1 (24-fold; p =0.0003) and BirA cultures (16-fold; p=0.001) and with more myeloid than erythroid cells (p=0.01). In all three genotypes ∼50-60% DN cells were Ter119^-^Gr1^-^Mac1^-^. Given FACS-purified DN cells showed higher mRNA expression of erythroid genes and lower expression of myeloid and megakaryocytic genes (**Figure 3C**), one possible lineage assignment for the Ter119^-^Gr1^-^Mac1^-^ cells could be immature Ter119^-^ erythroid cells.

**Figure 3:**
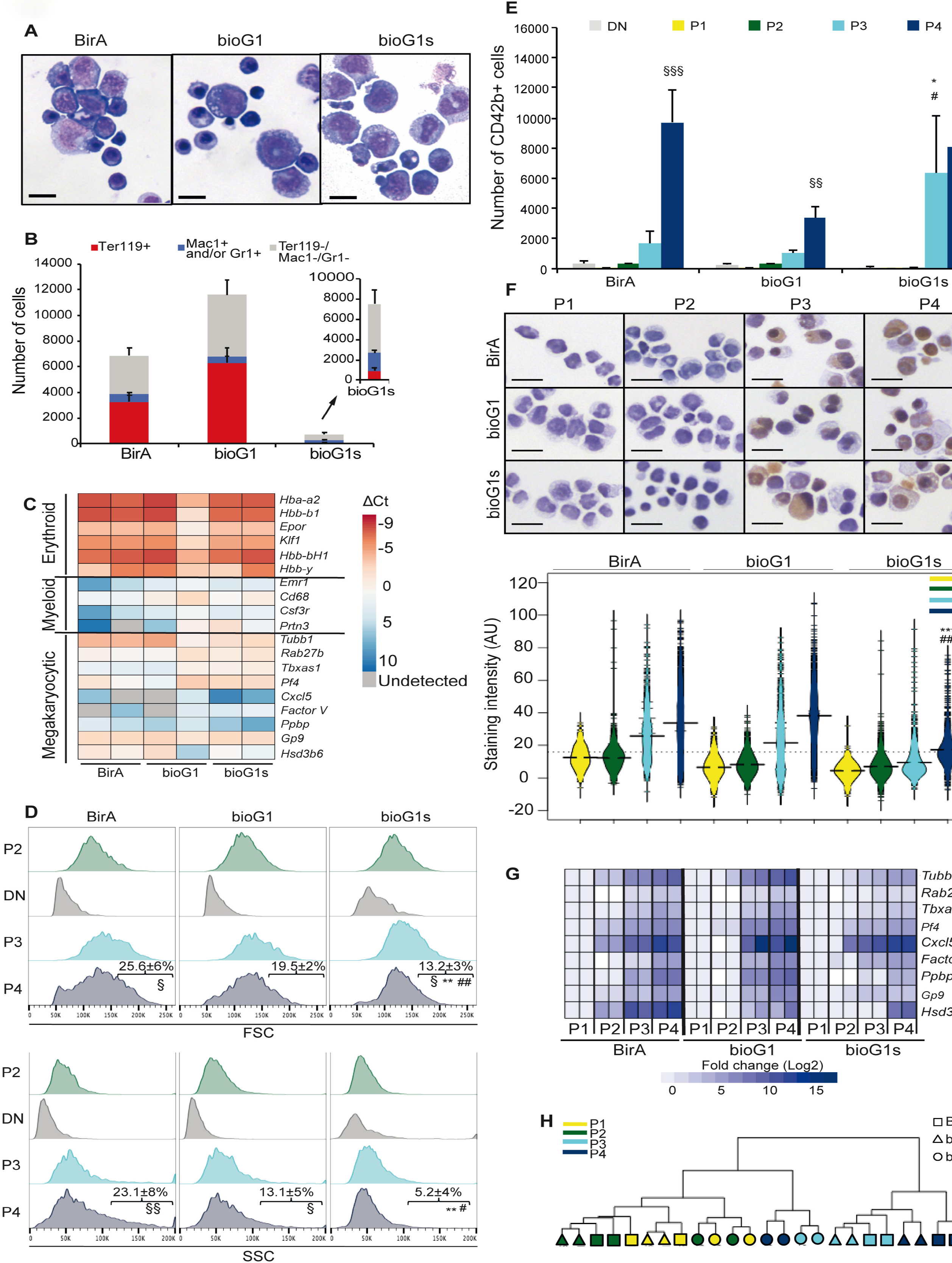
Gata1s cells fail to produce erythroid cells and mature megakaryocytes. A) Micrographs of May-Grunwald-Giemsa-stained cytospins of FACS-purified DN (kit^-^ CD41^-^) cells at day 10. Genotype of cells is indicated above. Scale bars represent 25μm. B) Bar plot of number of erythroid cells (Ter119^+^) and myeloid cells (Gr1^+^ and/or Mac1^+^) within the DN population at d10 of culture in BirA, bioG1 and bioG1s cultures. 250000 total cells analysed by FACS in each case. C) Heatmap of mRNA expression of selected erythroid (top), myeloid (middle) and megakaryocytic (bottom) genes (in rows) in DN BirA (left), bioG1 (middle) and bioG1 (right) cells at d10. Data from two independent biological replicates is shown. D) Representative histogram (three independent experiments were performed) showing size (FSC-A, top panel) and granularity (SSC-A, bottom panel) assessed by flow cytometry. Data from BirA (left), bioG1 (middle), bioG1s (right) from P2, DN, P3, P4 populations is shown. In P4, numbers indicate the mean percentage ±SD of cells within the gate. ^§^p<0.05 and ^§§^p<0.01 between P4 and P2. **p<0.01 between bioG1s and BirA. ^#^p<0.05 and ^##^p<0.01 between bioG1s and bioG1. FSC-A P2 vs P4: BirA p=0.03; bioG1 p=0.1; bioG1s p=0.02 SSC-A P2 vs P4: BirA p=0.003; bioG1 p=0.01; bioG1s p=0.06 P4 FSC-A>150K: BirA vs bioG1s p=0.0012; bioG1 vs bioG1s p=0.0059 P4 SSC-A: BirA vs bioG1s p=0.003, bioG1 vs bioG1s p=0.011 E) Bar plot showing the number of cells expressing CD42b at d12 from DN, P1, P2, P3 and P4 populations from genotypes indicated. ^§§^p<0.01 and ^§§§^p<0.001 between P4 and P3. *p<0.05 between bioG1s and BirA. ^#^p<0.05 between bioG1s and bioG1. BirA P3 vs P4 p=0.0002, bioG1 P3 vs P4 p=0.003. P3 CD42b+ cells: BirA vs bioG1s p=0.02. P3 CD42b^+^ cells: bioG1 vs bioG1s p=0.01 F) Top, representative micrographs of acetylcholinesterase (AChE) staining of FACS-purified P1 to P4 d10 BirA, bioG1 and bioG1s populations. Scale bars represent 10μm. Bottom, quantitation of AChE staining, from 500 cells, analyzed from three independent experiments. Bean-plot of staining intensity expressed in arbitrary units (AU) (y-axis) for each population (P1 to P4) from BirA, bioG1 and bioG1s cells. ***p<0.001 between bioG1s and BirA. ^###^p<0.001 between bioG1s and bioG1. P4: BirA vs bioG1s p=2.0E-130; bioG1 vs bioG1s p=7.7E-231. G) Heatmap of mRNA expression of selected megakaryocytic genes (indicated on the right) FACS-purified P1, P2, P3 and P4 at d10 (columns). Data from two independent biological replicates is shown. Genotype of the cells is indicated below the heatmap. H) Hierarchical clustering using mRNA data from (G).

### Altered megakaryocytic maturation of bioGATA1s cells

During differentiation, megakaryocytes enlarge considerably, acquire granules and develop a demarcation membrane system for proplatelet formation. We used multiple approaches to study megakaryocyte maturation as cells progressed from P2 to P4. First, morphologically, cells in populations P1 and P2 were small, with a blast morphology (**Supplemental Figure 5A**). In contrast, cells in P3 and P4 were larger and particularly in P4 were maturing megakaryocytes. To quantify these changes, we measured cell size (forward scatter, FSC-A) and granularity (side scatter, SSC-A) by flow cytometry (**Figure 3D**). Concordantly, there was a progressive increase of size and granularity from P2 to P3 and P4 (BirA: median FSC-A P2 120K±3 vs P4 129K±4 p<0.01; bioG1: median FSC-A P2 118K±4 vs P4 122K±4 p<0.1) (BirA: median SSC-A P2 59K±9 vs 81K±9 P4, p<10^−3^; bioG1: median SSC-A P2 59K+/-6 vs P4 69K±5 p<0.01). A similar trend was also seen in the mutant cells, but to a lower extent (P2 vs P4, FSC 110K±6 vs 124K±2 p<0.01, SSC 49K±3 vs 60K±8 p<0.01). Closer inspection of FSC and SSC profiles showed a lower proportion of larger and more granular cells in the P4 population in bioG1s compared to control BirA and bioG1 populations (FSC-A>150K: bioG1s, 13.2±3%; BirA, 25.6±6%; bioG1, 19.5±2%. bioG1s vs BirA p=0.0012, bioG1s vs bioG1 p=0.0059) (SSC-A>110K: bioG1s, 5.2±4%; BirA, 23.1±8%; bioG1, 13.5±6%; bioG1s vs BirA p=0.003; bioG1s vs bioG1 p=0.011). Finally, as expected the erythroid-dominant DN cells showed decreased cell size and granularity compared to P2 cells.

Next, we studied CD42b expression in P1 to P4 populations at day 12 (**Figure 3E, Supplemental Figure 5B-E**). As expected, very few cells in DN, P1 and P2 expressed CD42b (<4%; absolute numbers <200 cells). In contrast, and as expected, the absolute number (**Figure 3E**) and proportion (**Supplemental Figure 5E**) of CD42b^+^ cells in P3 and P4 were much significantly higher than in DN, P1 and P2. Importantly, there were significant differences between bioG1s and BirA and bioG1. Absolute numbers of CD42b^+^ in P3 (**Figure 3E**) were significantly greater in bioG1s compared to BirA and bioG1 supporting the hypothesis that compared to wild type GATA1, GATA1s promotes proliferation of kit^lo^ immature megakaryocytes (P3). In contrast, the absolute numbers of mature kit^-^CD42b^+^ P4 bioG1s cells were no different compared to bioG1 and BirA.

Furthermore, the absolute number and ratio of CD42b^+^ cells in P4 relative to P3 was greater in BirA and bioG1. In contrast, in bioG1s the absolute number, and ratio, of CD42b^+^ cells was not significantly different between P3 and P4 (**Figure 3E** and **Supplemental Figure 5E**). This supports the hypothesis that GATA1s, compared to wild-type GATA1, is less effective at driving maturation of P3 megakaryocytic cells to P4 megakaryocytic cells.

Next, we measured activity of acetylcholinesterase, an enzyme whose activity increases with megakaryocyte maturation, by quantitating intensity of an acetylcholinesterase driven cytochemical reaction in purified P1, P2, P3 and P4 populations (**Figure 3F**). The intensity of acetylcholinesterase-induced cytochemical staining was low in P1 and P2 and increased in P3 cells, and increased further, in P4 cells. Importantly, there was significantly lower cytochemical staining in P3 and P4 in bioG1s compared to control BirA and bioG1 cells, which may reflect the smaller size of P3 and P4 cells bioG1s and/or difference in maturation state of bioG1s cells.

Finally, we tested mRNA expression of megakaryocyte specific genes in P1-P4 in all three genotypes (**Figure 3G**). There was reduced expression of megakaryocyte genes in P3 and P4 in bioG1s compared to BirA and bioG1 cells (*Tubb1, Factor V, Pbbp, Gp9* and *Hsd3b6*). Hierarchical clustering analysis confirmed that bioG1s P3 and P4 cells were transcriptionally more closely related to the more immature P1 and P2 cell populations than P3 and P4 from GATA1 wild-type cells (**Figure 3H**).

Taken together, these data confirm megakaryocytes mature from P1 to P4. BioG1s produce more immature megakaryocytes (P3) but they fail to differentiate as efficiently into the most mature megakaryocyte population (P4) compared to wild type cells.

### Decreased apoptosis and increased proliferation in mutant P3 population

To understand why the number of bioG1s cells increased during megakaryocytic differentiation (**Figure 1G** and **Figure 3E**), we asked if this was due to reduced apoptosis (**Figure 4A-B**) and/or increased cycling of cells (**Figure 4C-D)** in bioG1s compared control BirA cells. We performed flow analyses in P1-P4 sub-populations at d8 (for apoptosis) or d9 (for cell cycle) (2 and 3 days after re-culture of kit^hi^CD41^lo^). There was a significant, and specific, decrease in Annexin V^+^ cells in bioG1s P3 cells (bioG1s, 4.77±0.75%; BirA, 10.57±1.33%; p=0.002). There was also a specific, and significant, increase of bioG1s P3 cells in S phase (bioG1s, 43.9±7.18%; BirA, 27.57±4.84%; p=0.007) and decrease in G1/G0 phase (bioG1s, 25.53±7.82%; BirA, 39.7±4.77%; p=0.03).

**Figure 4:**
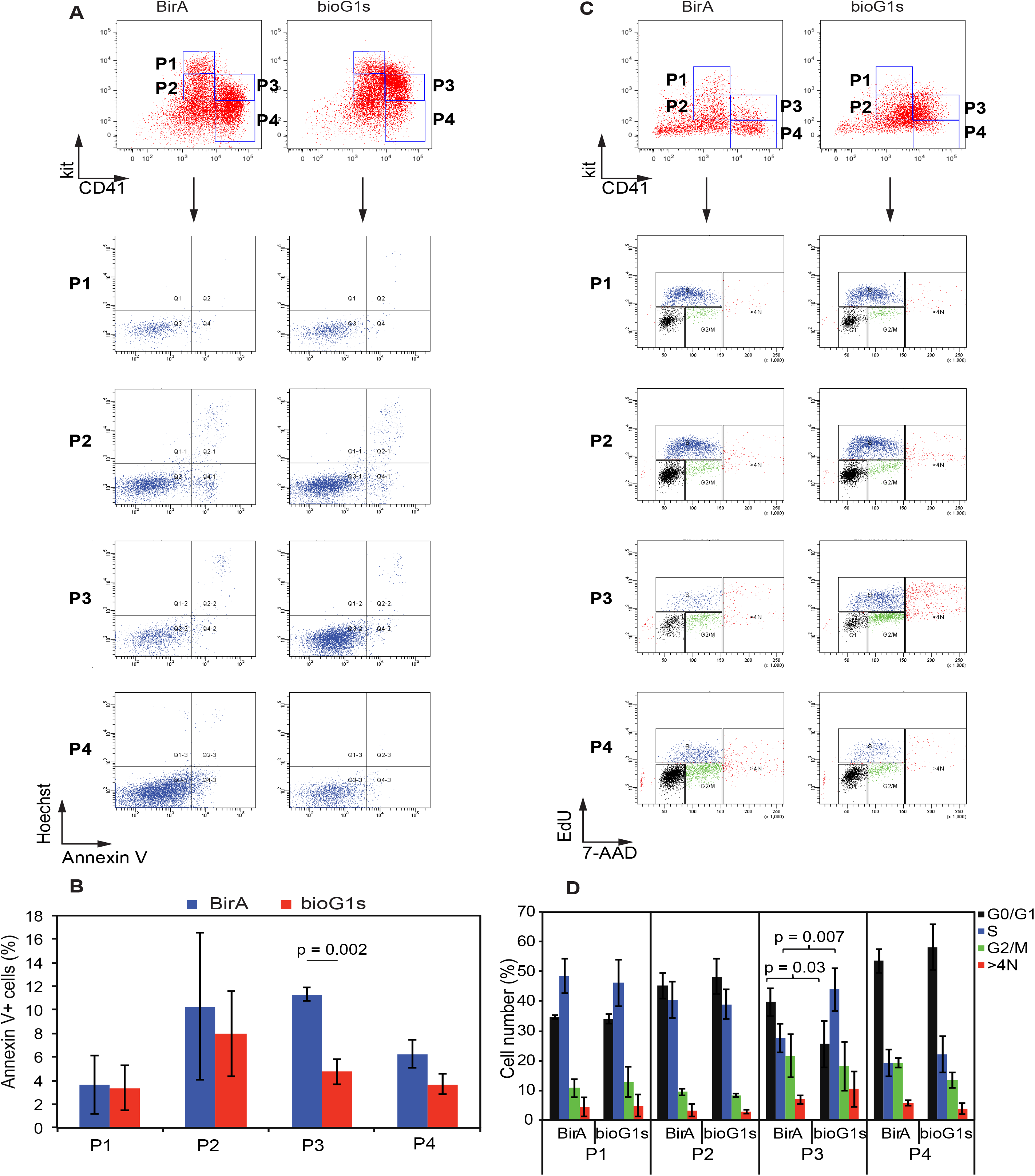
Increased proliferation and decreased apoptosis in GATA1s P3 cells. A) Flow cytometry analysis, from one representative experiment (of three independent experiments) at d8, showing kit and CD41 expression of BirA (left) and bioG1s (right) cells (top). P1 to P4 populations indicated. Below, Annexin V and Hoechst staining within P1 to P4 populations. B) Bar plot of data from all three experiments showing mean percentage ±1SD of AnnexinV^+^ cells in BirA and bioG1s cultures. P3 cells: p=0.002 C) Representative flow cytometry analysis, from one experiment (of three independent experiments) at d9. Details as set out in (A). Below, cell cycle analysis determined by EdU incorporation and 7-AAD staining. D) Bar plot from all three experiments showing mean percentage ±1SD of cells in G0/G1, S, G2/M phases of cell cycle and cells with >4N ploidy, in BirA and bioG1s cultures. P3 G0/G1: p=0.03; P3 S: p=0.007

### Modelling transitions through differentiation

Using the kinetic data of differentiation (**Figure 2**), together with absolute cell numbers produced per initial cell numbers and the cell cycle and apoptosis data (**Figure 4**) we have developed a mathematical model (see Supplementary Methods) to study the rates at which the cells transition between P1 to all other populations and how *Gata1s* mutation alters the kinetics of transition (**Figure 5**). The fit of the model to the data can be seen in **Figure 5A** and the modelled cell numbers in culture (**Figure 5B**) closely mirrors the actual cell numbers produced in culture (**Figure 1G**). Comparing the rates of transition between the different populations, for BirA and bioG1s cells, only the rate of transition of P3 and P4 was different between BirA and bioG1s cells. Here, bioG1s showed statistically markedly reduced transition between P3 and P4 compared to BirA cells. This slower transition from P3 to P4 produces an accumulation of cells in P3, where cells are proliferating more than in P4. This provides a likely explanation for the large increase in cell numbers seen in **Figure 1G** for bioG1s cells from D10 to D12.

**Figure 5:**
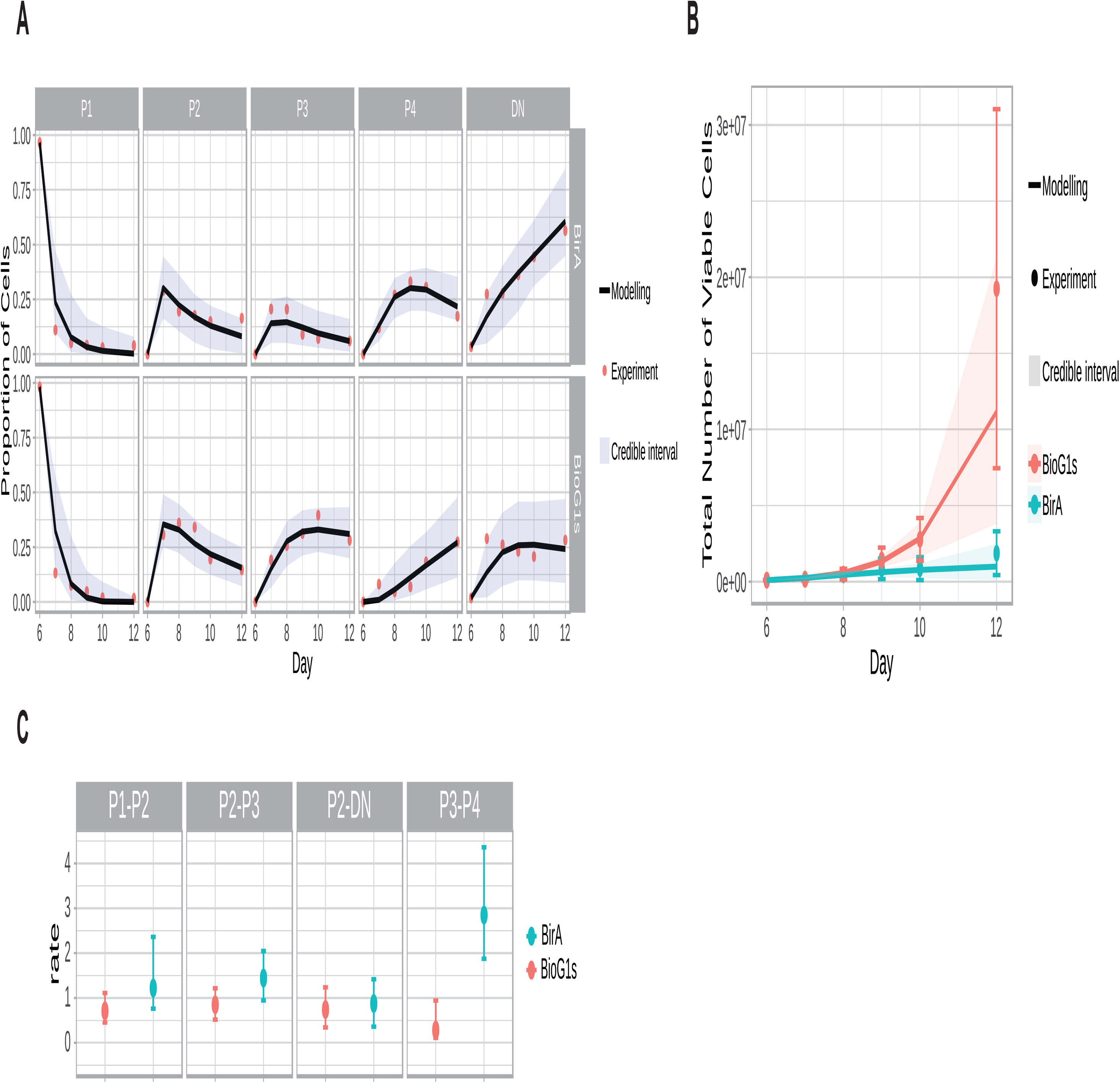
Cell fate modelling suggests GATA1s enhances cell division in P3 cells and reduces commitment to P4. A) Fit of the mathematical model to the proportions of cells in each population. Top, BirA cells, blow bioG1s cells. The points are the data taken from Figure 2 and the solid line is the model fit. The shaded region is the 95% credible interval for the data – i.e. 95% of the data should lie within the shaded region. B) Growth curve for the total number of cells modelled from the model fit. Top, BirA cells, blow bioG1S cells. Note the difference in scale of the y-axis. The error bars of the data are two times the standard deviation of the replicates. Note the rate of transition of P3 to P4 is much lower for bioG1s. C) Inferred cell transition rates between the populations and their 95% credible interval.

### GATA1s phenotype is recapitulated in vivo

Next, we asked if the *in vitro* EB-derived P1-P4 populations were present in mouse development. EB hemopoiesis mimics yolk sac hemopoiesis^17,18^ in that it first produces kit^+^CD41^+^CD16^-^CD32^-^ primitive yolk sac erythroid progenitors with myeloid and megakaryocytic potential^17,19^, followed by kit^+^CD41^+^CD16^+^32^+^ definitive erythroid-myeloid progenitors (EMPs)^20^. In d6 EB cultures, the majority of kit^hi^CD41^lo^ hemopoietic cells were CD16^-^CD32^-^ primitive erythroid progenitors with myeloid potential (**Figure 6A** and **Supplemental Figure 6A-C**) consistent with previous reports^20^.

**Figure 6:**
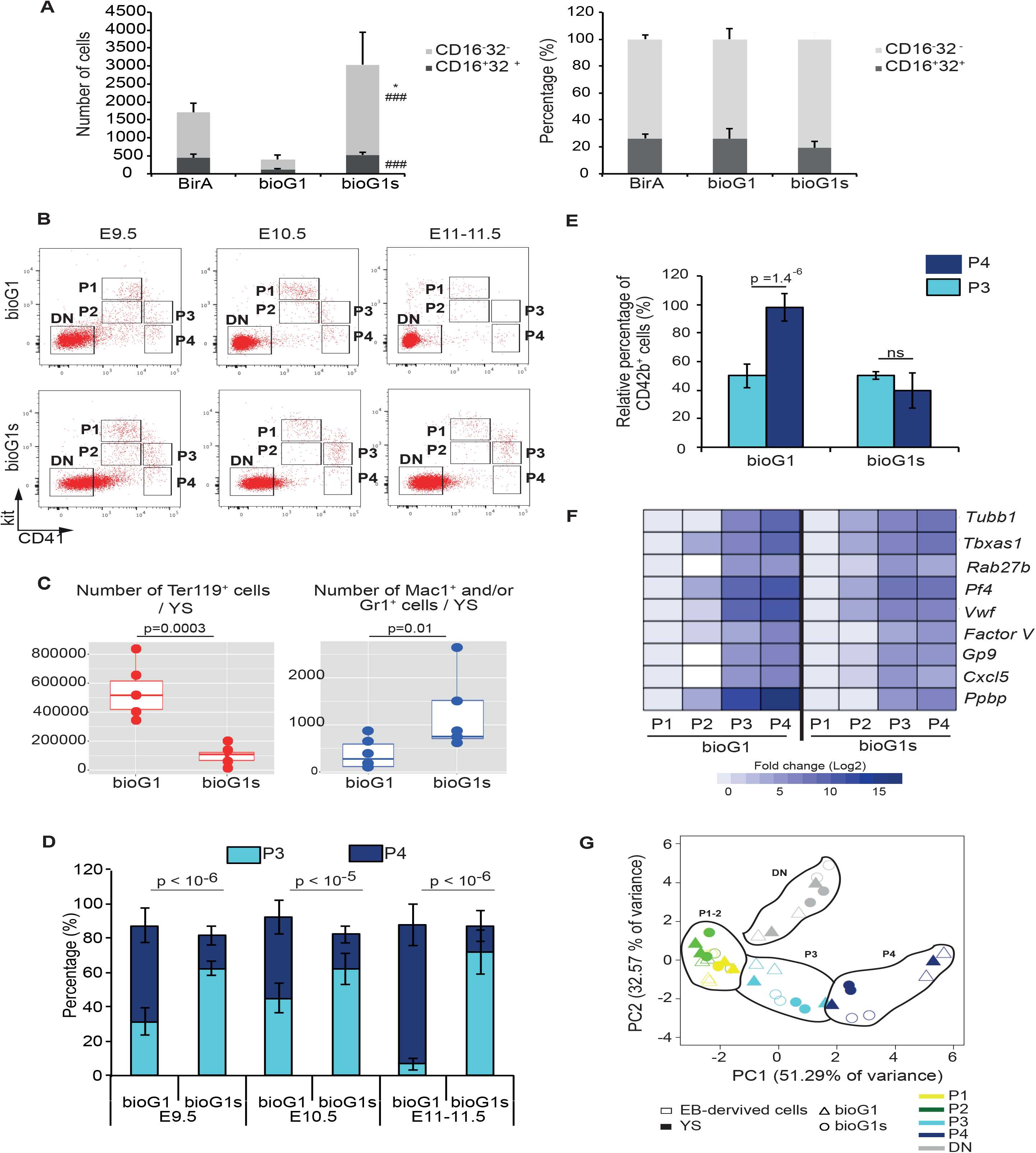
Hemopoietic populations in the yolk-sac. Expansion of GATA1s P3 relative to P4 populations. A) Bar plot of absolute number (left) and percentage (right) of primitive progenitors with myeloid potential (CD16^-^CD32^-^) and definitive erythro-myeloid progenitors (CD16^+^CD32^+^) in d6 EB-derived kit^hi^CD41^lo^ cells. Cell genotype is indicated. *p<0.05 versus BirA. ^###^p<0.001 versus bioG1. CD16/32- cells: BirA vs bioG1s p-value =0.017; bioG1 vs bioG1s p-value = 0.00064 CD16/32+ cells: BirA vs bioG1s p-value =0.075; bioG1 vs bioG1s p-value = 9.67E-6 B) Representative flow cytometry analysis plot of kit and CD41 expression from bioG1 (top) and bioG1s (bottom) at E9.5 (left), E10.5 (middle) and E11-11.5 (right) (n=5 to 7 yolk sacs analysed individually for each genotype). P1-P4 and DN populations indicated. C) Box plot of absolute number/yolk sac of erythroid (left, Ter-119^+^) and myeloid (right, Mac1^+^ and/or Gr1^+^) in bioG1 and bioG1s. Each dot represents one yolk sac analysed at E10.5 (n=5/genotype). Erythroid cells: bioG1 vs bioG1s p=0.003; myeloid cells bioG1 vs bioG1s p=0.01. D) Percentage of yolk sac cells (y-axis) within P3 and P4 compartments calculated from data shown in B for both genotypes (bioG1, bioG1s) at different timepoints (x-axis). E) Mean percentage ±1 SD of CD42b^+^ E10.5 yolk sac cells in P3 and P4 in bioG1 and bioG1s (n=5 for each genotype). bioG1: P3 vs P4 p=1.4E-6. F) Heatmap of fold change of mRNA expression of megakaryocytic genes (rows) in E10.5 yolk sac cells purified from P1 to P4 from bioG1 (left) and bioG1s (right). Cells purified from two independent litters for each genotype. G) 2D-PCA plot of mRNA expression of all genes from either YS cells (shaded symbols) or from EB-derived *in vitro* cultures (open symbols), from bioG1 (triangles) and bioG1s (circles) genotypes, from P1 to P4 and DN cells (color coded as indicated below the figure). Data taken from Supplemental Figures 5F (EB-derived cultures) and 6H (yolk sac).

Next, we analyzed hemopoietic cells from E9.5 to E11.5 yolk sac for kit and CD41 expression (**Figure 6B, Supplemental Figure 6D-F**). In E9.5 yolk sac, we identified populations with the same immunophenotypic profile as P1 to P4 and DN (kit^-^CD41^-^) populations seen *in vitro* in cultures, in both bioG1 and bioG1s embryos. From E9.5 to E11.5, the DN population was sustained in both bioG1 and bioG1s yolk sac. We then purified this population and quantitated the absolute number of cells/yolk sac (**Figure 6C** and **Supplementary Figure 6D-F**). Absolute numbers of DN cells were far lower in bioG1s yolk sac, with a significantly marked reduction in Ter-119^+^ cells and increase in Mac-1^+^/Gr1^+^ cells consistent with data from EB-derived cultures.

Turning to P1-P4 populations, there was a reduction of P1-P2 populations at E10.5, which virtually disappeared by E11-11.5 with mainly P3 and P4 populations present. Importantly, in bioG1s, there was a significant increase in P3 relative to P4 at each time point (E9.5-E11.5) and sustained higher levels of P3 cells at E11-11.5, mirroring *in vitro* culture data (**Figure 6B-D, Supplemental Figure 6D-F**). Purified P3 and P4 populations contained CD42b^+^ cells (**Figure 6E, Supplemental Figure 6D-G**). In control bioG1 cells there were significantly more CD42b^+^ cells in P4 than P3, consistent with megakaryocyte maturation in P4. This was not the case in bioG1s P4 cells, consistent with aberrant, reduced megakaryocyte maturation. We also tested mRNA expression in purified E10.5 P1-P4 cells (**Figure 6F**). In bioG1 cells megakaryocytic gene expression (most noticeable for *Ppbp, Vwf, Pf4, Tbxas1*) increased progressively from P1/P2 to P3 then to P4. In contrast, in bioG1s cells expression of these genes did not increase from P3 to P4 cells, consistent with a megakaryocyte maturation defect. Finally, we compared mRNA expression in yolk sac and EB-derived DN and P1-P4 populations using two-dimensional principal component analysis (PCA) (**Figure 6G and Supplemental Figures 5F and 6H**). The most important finding was that yolk-sac and EB-derived DN and P1-P4 populations clustered together, consistent with the notion that the yolk sac and EB populations are transcriptional similar. Principal component 1 (PC1) (51% of variance) separated the P3 and P4 populations (genes whose expression contributed most to variance were the megakaryocyte genes – *Tubb1, Factor V, Gp9* and *Pf4*) whereas PC2 (32% of variance) separated the P1-P2 and DN populations (genes whose expression contributed most to variance were the erythroid genes – *Klf1, Epor*, and *globin genes*).

## Discussion

Our studies of a new knock-in GATA1s allele, in hemopoiesis from ES cells and in murine yolk sacs, define the cellular mechanisms leading to a developmental-stage specific megakaryocyte myeloproliferation that likely contributes to the oncogenic effect of GATA1s. GATA1s results in a 10-fold increase in megakaryocytic cells from ES cultures compared to control. Though prior work on GATA1s TMD-derived induced pluripotent stem cells (iPSC) also demonstrated erythroid differentiation arrest and enhanced megakaryocyte differentiation, the stage in hemopoiesis where perturbed differentiation occurs was unclear^13^. We now demonstrate that accumulation of megakaryocytic lineage cells occurs predominantly late in megakaryopoiesis, at an immature megakaryocyte precursor stage (where most cells are 2N), within a specific compartment (termed P3), characterised by high CD41 expression and low level of kit expression (kit^lo^CD41^hi^).

A number of mechanisms may contribute to increased number of GATA1s P3 cells. Hemopoietic cells have a range of cell fate options including differentiation (with or without entering cell cycle), entering cell cycle without differentiating, apoptosis and quiescence. Our data shows that GATA1s P3 cells have increased number of cells in S-phase, reduced number in G0/G1 and a lower number of apoptotic cells compared to GATA1 P3 cells. Detailed kinetic studies of ES-derived hemopoiesis, demonstrate a delay in exiting the P3 compartment into the next, more mature megakaryocyte compartment (termed P4) where cells have lost kit expression and presumably lost the proliferative drive afforded by kit signalling. For a 10-fold increase in cell number there need only be just over 3 more cell divisions to account for the increase in GATA1s cell numbers.

Three major, open questions arise out of our work that provide a platform for future studies. The first two related questions are, what molecular mechanisms explain how GATA1s causes differentiation delay and why does differentiation delay specifically occur in the megakaryocyte lineage? Though the answers to these questions are unclear, prior data suggests that sustained elevated expression of GATA2 in GATA1s cells may play a role^21^. Chromatin occupancy by GATA2/E-box proteins/LMO2/FLI1/ERG/RUNX1 heralds megakaryocyte lineage priming and sustained GATA2 repression of specific loci is correlated with terminal megakaryocyte maturation^22^ and indirectly modulates megakaryocyte cell progression in GATA1 deficient megakaryocytes^23^. However, proof that GATA2 is pivotal for GATA1s oncogenicity is still required and if GATA2 is needed, the mechanism by which it delays megakaryocyte differentiation in GATA1s cells requires further work.

Prior work has also suggested that GATA1 may directly interface with cell cycle^24^. Consistent with this one report has shown that GATA1 directly binds pRB/E2F2 via amino acid residues in the N-terminal GATA1 domain that is deleted in GATA1s^25^. Normally, GATA1/pRB/E2F2 restrain uncommitted murine hemopoietic cell proliferation whereas GATA1s fails to bind pRB/E2F2 and fails to do this.

Our data also confirm that erythroid maturation is reduced in GATA1s cells consistent with prior work^13^. One mechanism for this may be reduced GATA1s binding to cis-elements of erythroid genes which was demonstrated in an erythroid-megakaryocyte cell line model^26^ that may cause a failure of terminal erythroid maturation, resulting in activation of an apoptotic program which is normally forestalled by GATA1 and erythropoietin^27^.

The third question is why does GATA1s exert a developmental-stage specific myeloproliferative effect? One possible explanation is that fetal liver-restricted IGF-1 signalling promotes E2F-induced erythro-megakaryocyte proliferation and that the extent of this proliferation is restrained by GATA1, but not GATA1s^28^. Additionally, post-natal bone marrow-specific type 1 interferon signalling may actively suppress GATA1s megakaryocyte-erythroid progenitor growth promoting resolution of TMD in the post-natal period^29^.

In summary, our work now establishes the stage to test the role of previously identified molecular players (GATA1s, GATA2, E2F proteins, pRB, IGF-1 and interferon signalling) and possibly new determinants that regulate transition into and out of P3-like compartment in vivo and regulate the commitment of P2-like cells into either megakaryocytic or non-megakaryocytic paths of differentiation.

## Acknowledgements

P.V. and I.R. are supported by Bloodwise Specialist Programme Grant 13001 and by the NIHR Oxford Biomedical Centre Research Fund. P.V. and CP are supported by programme grants from the MRC Molecular Haematology Unit (MC_UU_12009/11). C.G. is supported by a Wellcome Trust Clinical Training Fellowship.

## Authorship Contributions

GJ, NS, QC, HC, KS, BS, CG, JD and BS performed experiments, analyzed the data. GJ, NS wrote the manuscript. QC, HC, KS, BS and CG contributed to editing the manuscript. EM, IR, CP, and PV designed the study, analyzed and interpreted the data, wrote the manuscript and academically drove the project.

