## Supplemental material for "Oncogenic *Gata1* causes stage-specific megakaryocyte differentiation delay"

#### Supplemental Information

#### Supplementary Methods

*Targeting constructs and targeting approach:* A murine *Gata1* gene DNA fragment (exon 1 to exon 5, 2.5 kb fragment upstream and a 2.1 kb downstream) was generated by PCR using 129/Ola strain DNA. For bioG1 targeting construct, the avi tag[1] was introduced in frame at exon 2 translational start codon via the *Nco I* restriction site. For bioG1s targeting construct, sequence including exon 2, intron 2 and the beginning of exon 3 (up to the translational start at amino acid 84) was excised and replaced with sequence coding an avi-TEV-FLAG peptide, resulting a fusion N-terminal GATA1s protein isoform with the avi-TEV-FLAG peptide. Targeting fragments were inserted either side of a Phosphoglycerate Kinase -neomycin (pgk-neo) cassette flanked by loxP sites. A negative selection cassette expressing the thymidine kinase gene regulated by MC1 promoter was included. IB10 ES cell line (subclone of E14) expressing a *BirA* transgene from the ROSA26 locus[2] was electroporated with targeting constructs and selected using G418 (0.3 mg/mL) and Ganciclovir (0.3 mM). Resistant colonies were characterized by PCR and Southern blotting. Normal karyotype ES clones (more than 80% of cells) were transiently electroporated with *Pgk-Cre* plasmid to delete the Neo cassette. Deletion was confirmed by PCR.

*Growth and differentiation of murine ESCs:* ESC lines were grown at 37°C in humidified 5% $CO_2$ incubator and maintained for up to 15 passages on gelatin-coated wells in KO-DMEM (Life Technologies, Warrington UK) supplemented with leukemia-inhibitory factor (LIF), 15% fetal calf serum (FCS), 2mM L-glutamine and 100uM β-mercaptoethanol. The protocol to differentiate megakaryocytes from ESC modified from a previously published report[3]. Briefly, one day prior to embryoid body (EB) differentiation, cells were passaged into IMDM (Life Technologies, UK) with LIF, FCS and 0.3 mM monothioglycecerol (MTG). $2x10^4$ to $3x10^4$ ESCs were plated into 100-mm Sterilin dishes in IMDM with 5% PFHMII, 15% FCS, 2 mM L-glutamine, Penicillin-Streptomycin, 0.4 mM MTG, 300 µg/ml human transferrin and 50 µg/ml ascorbic acid. After 6 days in culture, EBs were dissociated with 0.25% Trypsin/EDTA. Cells were incubated with PE-conjugated anti-mouse CD41. Antibody-labelled cells were isolated with anti-PE magnetic microbeads (Miltenyi, Bisley UK) and stained with APC-conjugated anti-mouse CD117 (kit). List of the antibodies used for the flow cytometry analyses is shown in Supplementary table 1. FACS purified CD41$^{+}$kit$^{+}$ cells were seeded onto OP9 cells with 20 ng/ml TPO (Peprotech, London UK). After 3 days, cells were seeded onto fresh OP9 cells with 10ng/ml TPO, 5ng/ml IL6 and IL11 (Peprotech, UK). Cells were maintained under these conditions for up to 6 days.

*Mice:* Animal studies were conducted with in accordance with the UK Home Office regulations. Embryos were staged based on somite pairs and morphological criteria. Embryos were removed from the uterine sac and kept on ice in DPBS (Life technologies UK) containing 10% FCS and 1% Penicillin-Streptomycin (Life Technologies, UK). Isolated yolk sacs were dissociated for 20 minutes at 37°C in PBS with 0.1% collagenase.

*Flow cytometry:* FACS analyses were performed either on LSR Fortessa X20 or FACS Canto II (BD, UK). FACS sorts were performed on FACS Aria Fusion (BD). Cells were labelled for 20 min at 4°C with Hoechst 33258 (Invitrogen, UK) and rat anti-mouse antibodies directed against: kit-APC, CD41-PE, Ter119-PECy7, Mac1-APCCy7, Gr1-FITC, and CD42b-FITC List of the antibodies used for the flow cytometry analyses is shown in Supplementary Table 1. Unstained, single stained and Fluorescence Minus One (FMO) controls were used to determine the background and fix the compensation in each channel. Data were analysed by FlowJo v10.1 software.

*GATA1 Western blotting:* This was performed as described[4] with modifications. Day 6 EB CD41$^{+}$ cells were lysed in hypotonic buffer containing 10 mM Hepes pH 7.9, 1.5 mM $MgCl_2$, 10 mM KCl (Sigma-Aldrich Poole UK) and a protease inhibitor cocktail (Roche Welwyn Garden City UK). Cell pellets were resuspended in hypertonic buffer (50 mM Hepes pH 7.9, 50 mM KCl, 300 mM NaCl, 0.1 mM EDTA, 10% Glycerol and a protease inhibitor cocktail). Sonicated lysates were incubated for 1h with 5 U/µl benzonase (Sigma-Aldrich). Protein concentration was determined using the Bradford Protein Assay (BioRad, Kidlington UK). 30 µg of nuclear extract were loaded onto 4 to 12% NUPAGE BisTris gradient SDS-PAGE gels

(Invitrogen, Paisley UK) and transferred onto a PVDF membrane. After blocking overnight at 4°C in 5% milk, the blots were incubated with anti-GATA1 (sc-1234 Santa Cruz Biotechnology, M20, 1/200) and TBP (sc-204 Santa Cruz Biotechnology, 1/500), and secondary HRP-conjugated antibodies. Blots were revealed by chemiluminescence (GE Healthcare Life Science, UK).

*Gene expression analysis by dynamic arrays*: Cells were FACS sorted into 9 µl of reaction buffer containing 5 µl of CellsDirect 2X reaction mix, 0.1 µl of RNase inhibitor, 2.5 µl of 0.2X Taqman gene expression assay mix, 1 µl of RT/Taq mix (all Life Technologies UK) and 0.4 µl of TE buffer. Reverse transcription and specific target amplification steps were performed as follows: 15 min at 50°C, 2 min at 95°C, 22 cycles of 15 sec at 95°C, 4 min at 60°C. cDNA diluted 1:5 in TE buffer was PCR-amplified with 48.48 dynamic array (Fluidigm, Cambourne UK). Each sample was analysed in technical duplicates or triplicates. Conditions with a quality score < 0.65 were considered as undetected. Biomark data were exported from the Fluidigm Data Collection Software and analysed using the $2^{-\Delta\Delta Ct}$ method. *GAPDH*, *TBP* and *HPRT* were used as housekeeping genes. For each gene, normalised ΔCt was calculated by subtracting the mean Ct of the three housekeeping genes to the gene of interest[5]. Downstream analyses were performed in R-3.3.3: pheatmap (heatmaps), hclust (hierarchical clustering with Euclidean distance measure and Ward.D2 agglomeration method), prcomp (PCA analysis using TRUE for the scale parameter), predict (for projection) and graphics (using the graphics package).

*Cell staining and microscopy*: Cells were cytospun for 5 min at 400 rpm and air dried overnight and stained with May-Giemsa-Grunwald (MGG) (Sigma, Welwyn Garden City UK) or acetylcholine iodide (Sigma) to reveal acetylcholinesterase activity. Pictures were taken using an Olympus BX60 microscope with an Infinity 3S Luminera color camera.

*Acetylcholinesterase staining quantitation:* All images were analysed by Fiji/ImageJ macroscript available at (https://github.com/dwaithe)[6]. Images were color inverted to create a negative image whereby the cellular staining was light against the background and the nuclear signals were particularly bright. Next, the image was smoothed lightly (sigma = 2.0) with a Gaussian kernel to reduce noise. The ImageJ ‘Find Maxima’ plugin was applied (noise tolerance = 16) to locate each cell nucleus and the ‘Maxima Within Tolerance’ option used to export a binary image of cell nuclei. Output was processed by “Analyze Particles” plugin excluding any objects < 50 pixels in size, considered too small to be nuclei. From a mask image of the retained regions a Voronoi transformation was applied to demarcate the position of the nuclei. In this way, it was possible to threshold (Otsu) the entire cell and accurately associate each nucleus with its corresponding cytoplasmic staining. The original image was then color deconvolved using the “H&E DAB” parameter set and the intensity in the resulting ‘Colour_1’ (blue) channel measured in the nuclear and cytoplasmic regions. Cytoplasmic and nuclear measurements were repeated for the ‘Colour_2’ (brown) deconvolved channel. After measurements, cellular masks were combined and selection inverted, and the background intensity measured in brown and blue channels. Data was then exported to Excel. For each picture in the brown channel the background intensity was subtracted to the cytoplasmic intensity. Beanplots were generated in R-3.3.3 using beanplot package.

*Cell cycle:* At day 9 of mESC differentiation, cell cycle was analysed with Click-iT EdU Alexa Fluor 488 Flow Cytometry Assay Kit (C10425; ThermoFisher Scientific, Hemel Hempstead UK). Cells were incubated for 1 h at 37°C with 10 μM EdU, then harvested and labelled with PE conjugated anti-kit and APC conjugated anti-CD41 antibodies for 20 minutes at 4°C. CD41/kit subpopulations were FACS-purified (FACS Aria II BD, UK). After fixation and permeabilization, Click-iT EdU Alexa Fluor 488 was detected and DNA was stained with SYTOX Advanced Dead Cell Stain Kit (S10349, ThermoFisher Scientific). Cells were analysed on a FACS Canto II (BD Biosciences).

*Apoptosis*: At day 8, cells were harvested and labelled with APC-conjugated anti-kit and PE-labelled anti-CD41 antibodies for 20 minutes at 4°C, washed in PBS with 10% FCS, resuspended in Binding Buffer (556454; BD Biosciences) and labelled with FITC conjugated Annexin V (A13199, ThermoFisher Scientific, UK). DNA was stained with SYTOX Advanced

Dead Cell Stain Kit (S10349; ThermoFisher Scientific) at room temperature for 15 minutes.

Cells were analysed on a FACS Canto II (BD Biosciences).

*Mathemathical Modelling*: For the mathematical model we use a system of differential equations that describe the total number of cells for each population (p1, p2, p3, p4 and pDN) and how they change over time.

$$\dot{x}_i = \sum_j A_{ij} x_j$$

Most of the $A_{ij}$ are zero, as we only allow transitions described experimentally (see fig 2C from main text). Also, aside from the cell growth, all the rates are balanced so that they represent movement between compartments. The above equation can be solved using the matrix exponential form:

$$X(t) = e^{At} X_0$$

We use a Bayesian approach to infer the model parameters from the data. The data used for fitting are the time series of measured proportions. We convert the solution of the ODEs to proportions $\pi_i(t) = x_i(t) / \sum x_j(t)$ and use a beta likelihood for the measured proportions $p_i$

$$p_{ik} \sim Beta(\pi_i(t_k)\sigma_i, (1 - \pi_i(t_k))\sigma_i)$$

Where $\sigma_i$ is a population specific parameter that controls the variability of the distribution. The proportions are sufficient to infer the transitions, however information on total cell numbers is not constrained. To constrain this part of the model we can use the total cell numbers that were experimentally measured separately. Rather than constraining the total number to a single value we specify that it comes from a normal distribution with mean the empirical mean and variance the empirical variance.

For the priors on the transition parameters we used a half-nomal(0, 2).

The Bayesian model was coded using the probabilistic modelling language Stan

#### Supplementary Figure Legends

**Supplemental Figure 1. Targeting of murine Gata1 locus and flow cytometry gates for Figure 1.**

A) Top, murine *Gata1* (m*Gata1*) locus. Exons 1-6 are numbered (blue boxes). Below, targeting constructs used to generate the bio*GATA1* and bio*GATA1s* knock-in alleles with position of 5' and 3' homology arms (red boxes), sequence encoding the AVI-TEV-FLAG tag (yellow box), the *neomycin* resistance gene (green arrow) and the *loxP* sequences for cre-recombinase.

B) Positions of the restriction enzymes sites (H, *Hind*III and B, *Bst*XI) with sizes (in base pairs) of the DNA restriction fragments and DNA probes (A and B) used for Southern blot screening in the wild type, modified bioGATA1 and bioGATA1s loci. Loci are depicted as in (A).

C) Southern blot analysis of *Hind*III (top panel) and BstXI (bottom panel) digested ES cell DNA.

D) Position of forward (fwd) and reverse (rev) primers in m*Gata1* loci used for PCR genotyping. Sizes of the amplified fragments are shown.

E) Agarose gel of PCR amplified DNA products, using fwd and rev primers shown in D, using template ES DNA as indicated.

F) Representative flow cytometry plots showing the serial gating strategy to analyse cultures at day 6 (d6) (top) and day 12 (d12) (below) based on morphology, isolation of single cells and Hoechst exclusion. On the right, Fluorescence Minus One (FMO) controls used to set positive gates for flow cytometry plots in Figure 1E.

G) Representative flow cytometry plots of kit and CD41 expression of d6 BirA (left), bioG1 (middle) and bioG1s (right) cultures of before (top) and after (below) CD41 bead enrichment. The box in the plots indicates the cell population sorted for d6-d12 cultures in the experiments.

H) Representative flow cytometry plots showing the serial gating strategy for flow cytometry plots in Figure 1F. Top, gates based on morphology, isolation of single cells and Hoechst exclusion. Below, single stained (left) and unstained (right) controls.

**Supplemental Figure 2: Gating Strategy for Flow Cytometric Plots in Figure 2.**

A) Representative flow cytometry plots showing serial gating strategy to isolate live-singlet cells based on morphology and Hoechst exclusion. Right, Fluorescence Minus One (FMO) controls used to set positive gates for Figure 2D.

B) Gating strategy to isolate subpopulations from bioG1s and BirA clones as well as post-sort purity analysis for subpopulations sorted from bioG1s clone.

**Supplemental Figure 3: FACS analysis of re-cultured FACS-purified P1-P4 populations from BirA and bioG1s cultures.**

A) FACS-purified hemopoietic progenitors, $kit^{hi}/CD41^{lo}$ (P1) from d6 EBs were cultured on OP9 stromal cells with cytokines for another 6 days. Representative flow cytometry plots of kit and CD41 expression from BirA (left), bioG1 (middle) and bioG1s (right) cultures. Aliquots of culture were analyzed daily for kit and CD41 and were monitored for 6 additional days (d7 to d12). The numbers indicated in the gates represents the mean percentage (± SD) of parent population from 3 independent experiments.

B) Representative FACS plots of kit and CD41 expression. P1-P4 populations from either BirA or bioG1s at d8 were FACS-purified (left plot) and then re-cultured for another two days (d8+1, middle plot; d8+2, right plot). The numbers indicated in the gates represent the mean percentage (± SD) of parent population from 3 (BirA) to 4 (bioG1s) independent experiments.

**Supplemental Figure 4: Flow cytometric gating and post-sort purity check for P1, P2, P3 and P4 for Figure 3.**

A) Flow cytometry plots of Fluorescence Minus One (FMO) controls for kit, CD41, Ter-119, Gr1 and Mac1 used to set gates for experiments in Figure 3B.

B) Flow cytometry plot of gating strategy to isolate live-singlet cells based on morphology and Hoechst exclusion for Figure 3B.

C) Representative flow cytometry plots of DN ($kit^{-}CD41^{-}$) cells (left) and expression of Ter119 (middle), Gr1 and Mac1 (right) in DN cells – data in Figure 3B. Numbers in the gates are mean percentages (± SD) of the parent population from 3 independent experiments.

D) Representative flow cytometry plots showing gates used to sort each population at d10 for experiments in Figure 3A-H.

E) Representative flow cytometry plots showing post-sort purity of sorted populations.

F) Table summarising mean (± SD) percentage purity of each sorted population (mean of 2 independent experiments).

**Supplemental Figure 5: Megakaryocyte differentiation and maturation**

A) Photographs of May-Grunwald-Giemsa staining of purified P1, P2, P3 and P4 d10 cells from BirA (top), bioG1 (middle) and bioG1s (bottom). Scale bars, 10 μm. N = 3 experiments.

B) Representative flow cytometry plots showing Fluorescence Minus One (FMO) controls for experiment in Figure 3E.

C) Representative flow cytometry plots of gating strategy to isolate single cells based on morphology and Hoechst exclusion for experiment in Figure 3E.

D) Representative flow cytometry plots of CD42b expression in d12 DN and P1-P4 populations in Bir A (top), bioG1 (middle) and bioG1s (bottom). Left, populations were first gated on kit and CD41 expression and then analysed for CD42b (right). In CD42b plots mean percentage (± 1SD) of $CD42^+$ cells (relative to parental) population from 4 independent experiments is indicated.

E) Bar plot of mean (± 1SD) percentage of $CD42b^+$ cells in DN, P1, P2, P3 and P4 populations for each genotype. N = 4 independent experiments. p-values were calculated using Student t-test.

F) Heatmap of mRNA expression of erythroid, myeloid and megakaryocytic genes in d10 P1-P4 populations. Genes are displayed horizontally. Duplicate biological samples were analyzed (displayed vertically). Cell genotype is indicated below the heatmap.

**Supplemental Figure 6: *In vivo* validation of the GATA1s phenotype.**

A) Representative flow cytometry plots showing Fluorescence Minus One (FMO) used to set gates in the experiment shown in Figure 5A.

B) Representative flow cytometry plots showing gating strategy followed to isolate live-singlet cells based on morphology and Hoechst exclusion for the experiment shown in Figure 5A.

C) Representative flow cytometry plots of kit and CD41 expression (top) and CD16/CD32 expression (below) in d6 EB derived cells. Numbers show the mean percentage (± 1SD) of parent population from 3 independent experiments.

D) Representative flow cytometry plots showing Fluorescence Minus One (FMO) used to set positive gates in the experiments shown in Figure 5C and E.

E) Representative flow cytometry plots showing the gating strategy followed to isolate live-singlet cells based on morphology and Hoechst exclusion for the in the experiments shown in Figure 5C and E.

F) Representative flow cytometry plots of kit and CD41 expression from bioG1 (top) and bioG1s (bottom) E10.5 yolk sac cells. Expression of Ter119, Gr1 and Mac1 was analysed as indicated. These data have been used to generate the box plots shown in Figure 5C.

G) Representative flow cytometry plots of CD42b expression from bioG1 (top) and bioG1s (bottom) E10.5 yolk sac cells from P1 to P4 populations. These data have been used to generate the box plots shown in Figure 5E.

H) Heatmap of mRNA expression of megakaryocytic, erythroid and myeloid genes in bioG1 (left) and bioG1s (right) E10.5 yolk sac DN, P1-P4 populations. Genes are displayed horizontally. Two biological duplicate samples were analysed for each population.

#### Supplementary Tables

**Table S1**: **Table of all antibodies used**.

#### Supplemental Figure 1

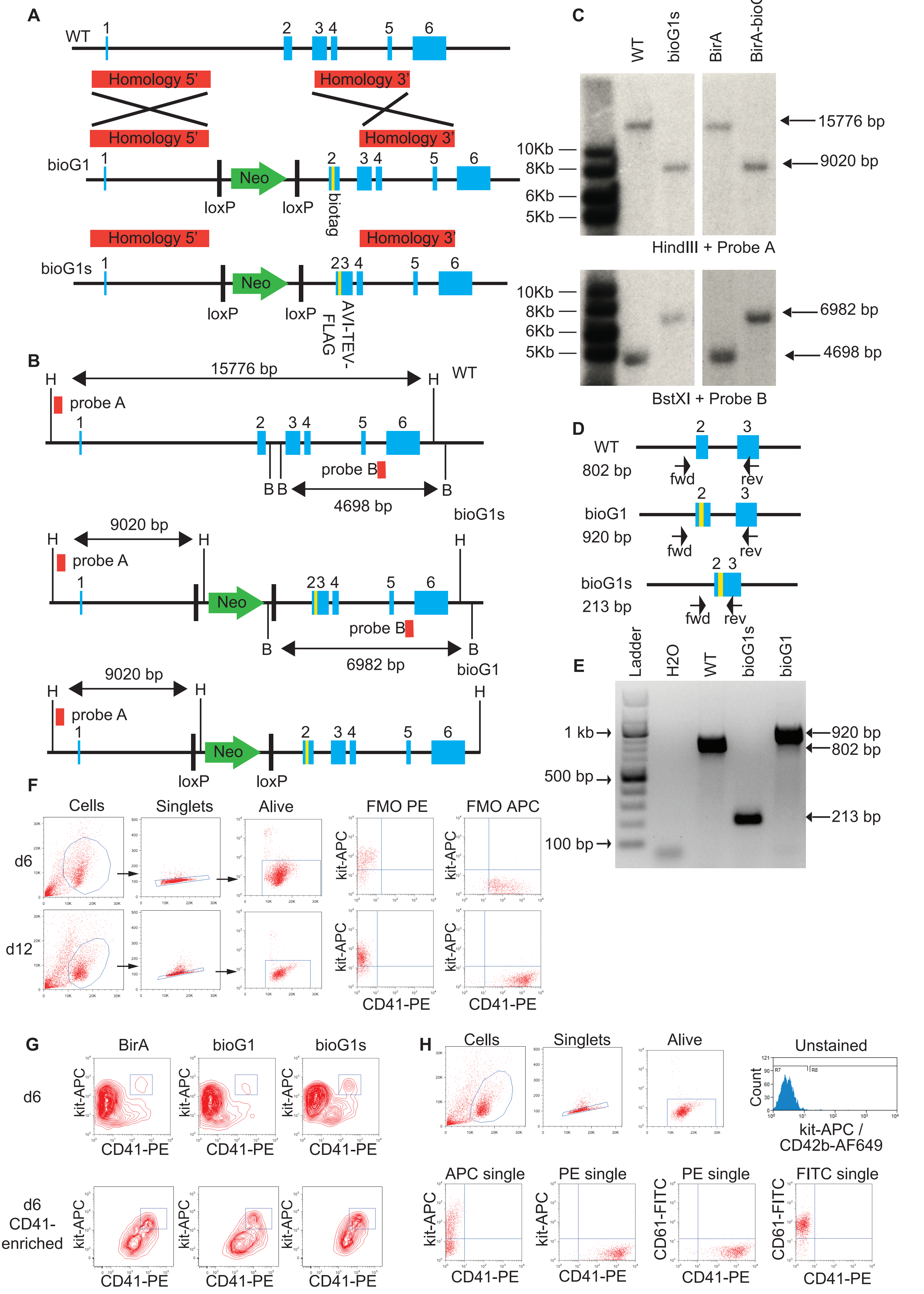

#### Supplemental Figure 2

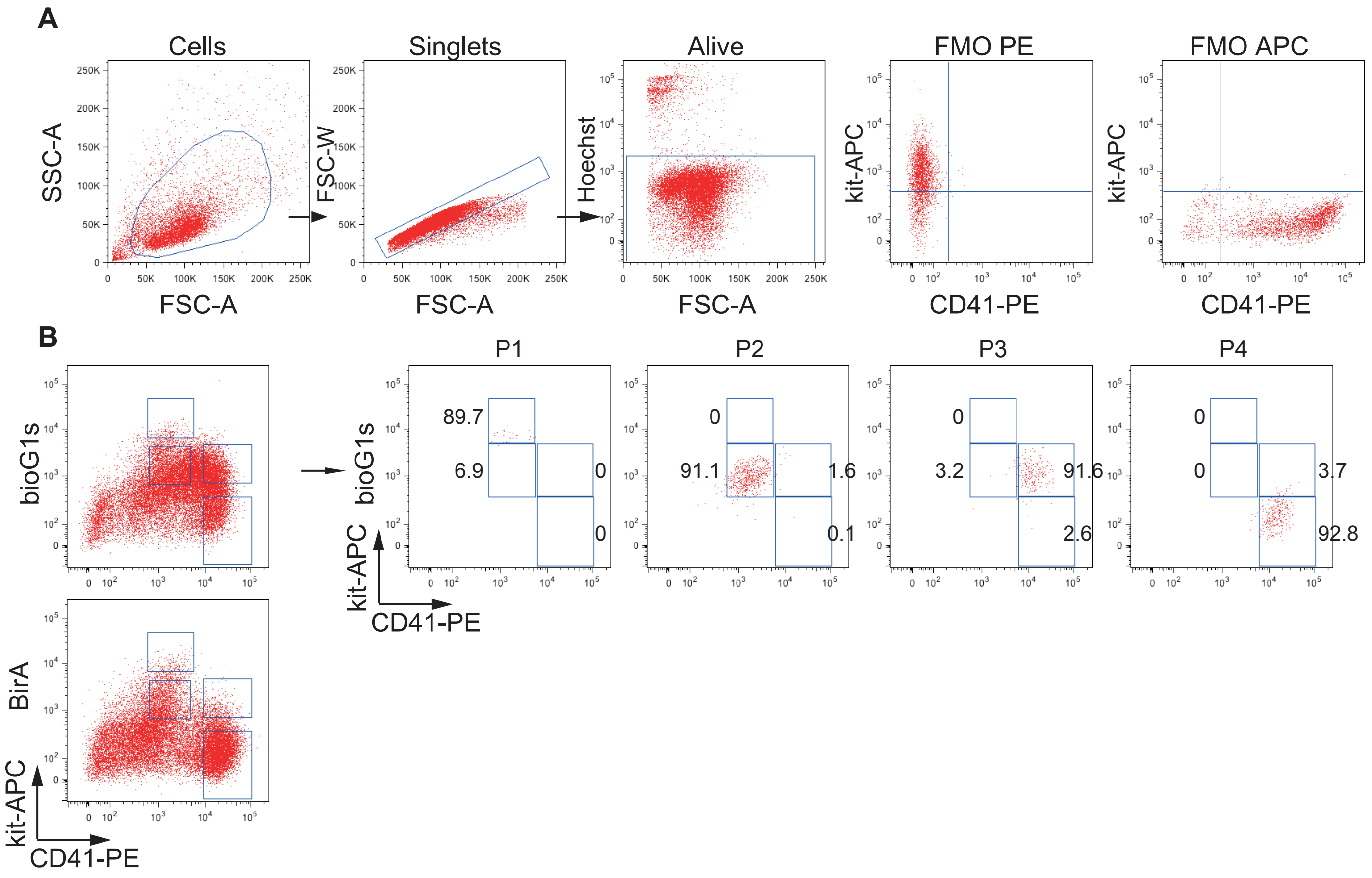

#### Supplemental Figure 3

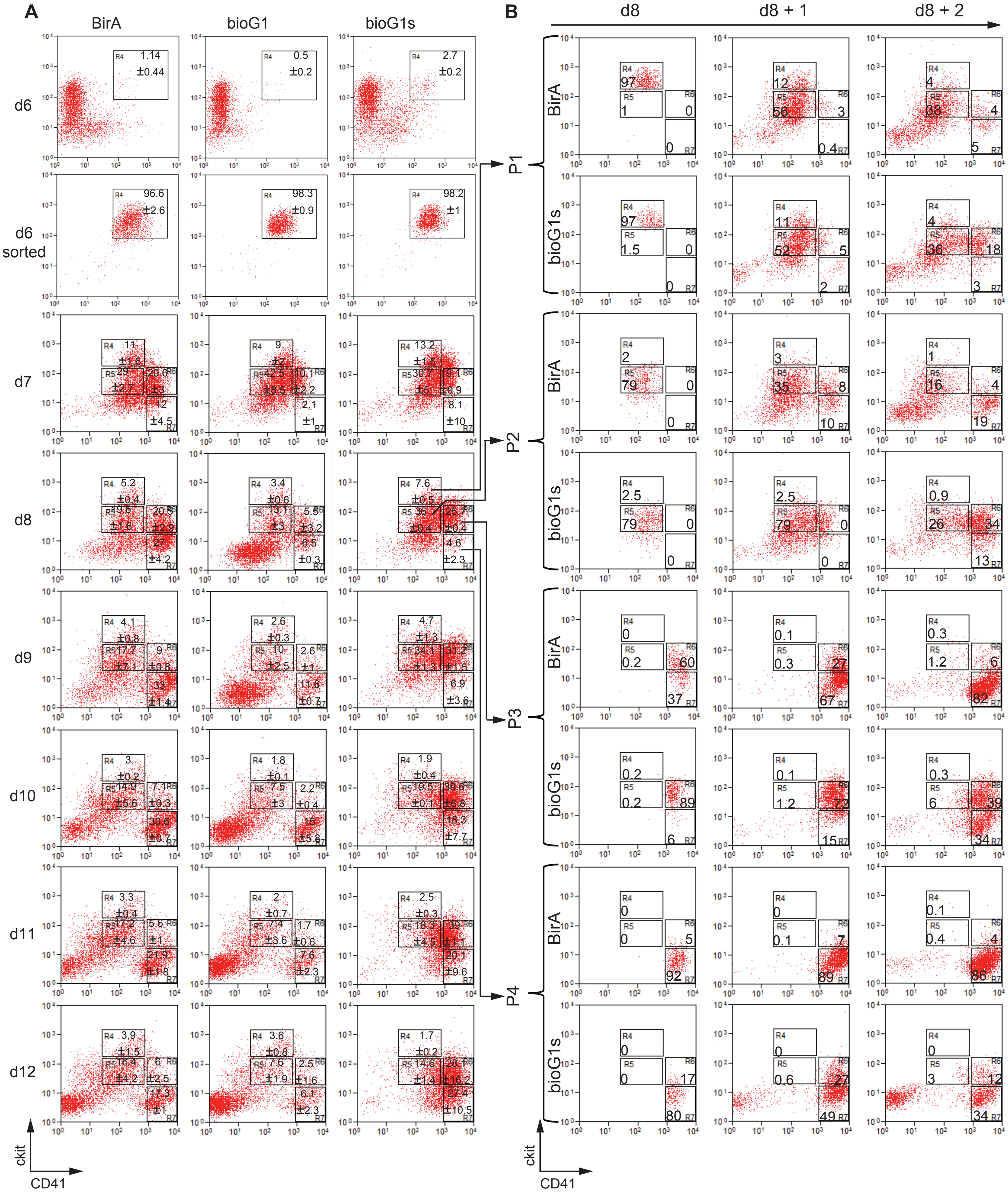

#### Supplemental Figure 4

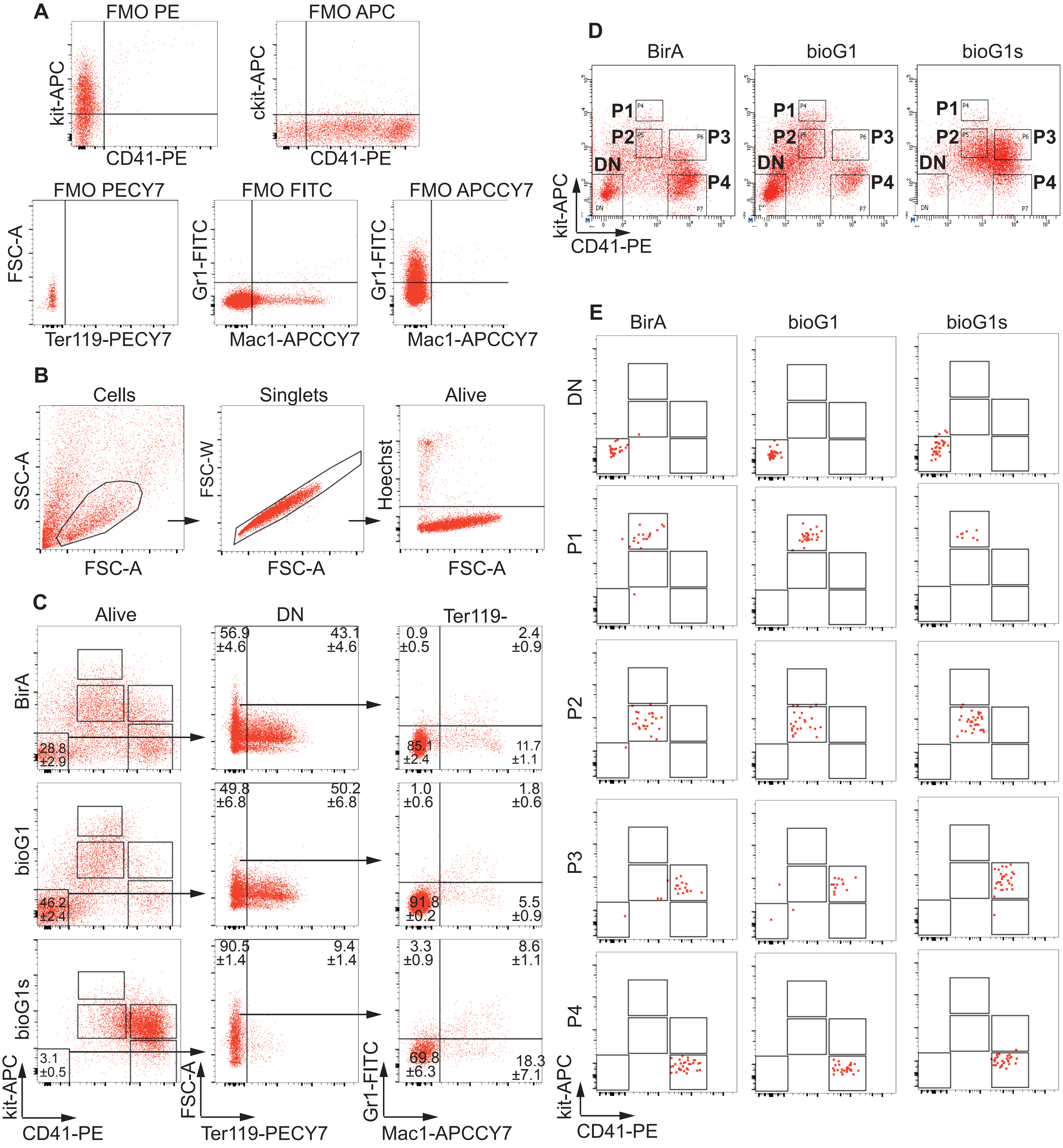


F

| | DN | P1 | P2 | P3 | P4 |
|---|---|---|---|---|---|
| BirA | 96.2±0.5 | 89.5±5.5 | 94.8±4.6 | 89.9±6.6 | 89.2±14.7 |
| bioG1 | 99.4±0.5 | 89.1±3.4 | 97.6±2.6 | 87.5±6.5 | 88.2±14.7 |
| bioG1s | 90.0±14.1 | 95.4+/-5.1 | 97.7±2.0 | 90.2±5.9 | 87.8±12.5 |

### Supplemental Figure 5

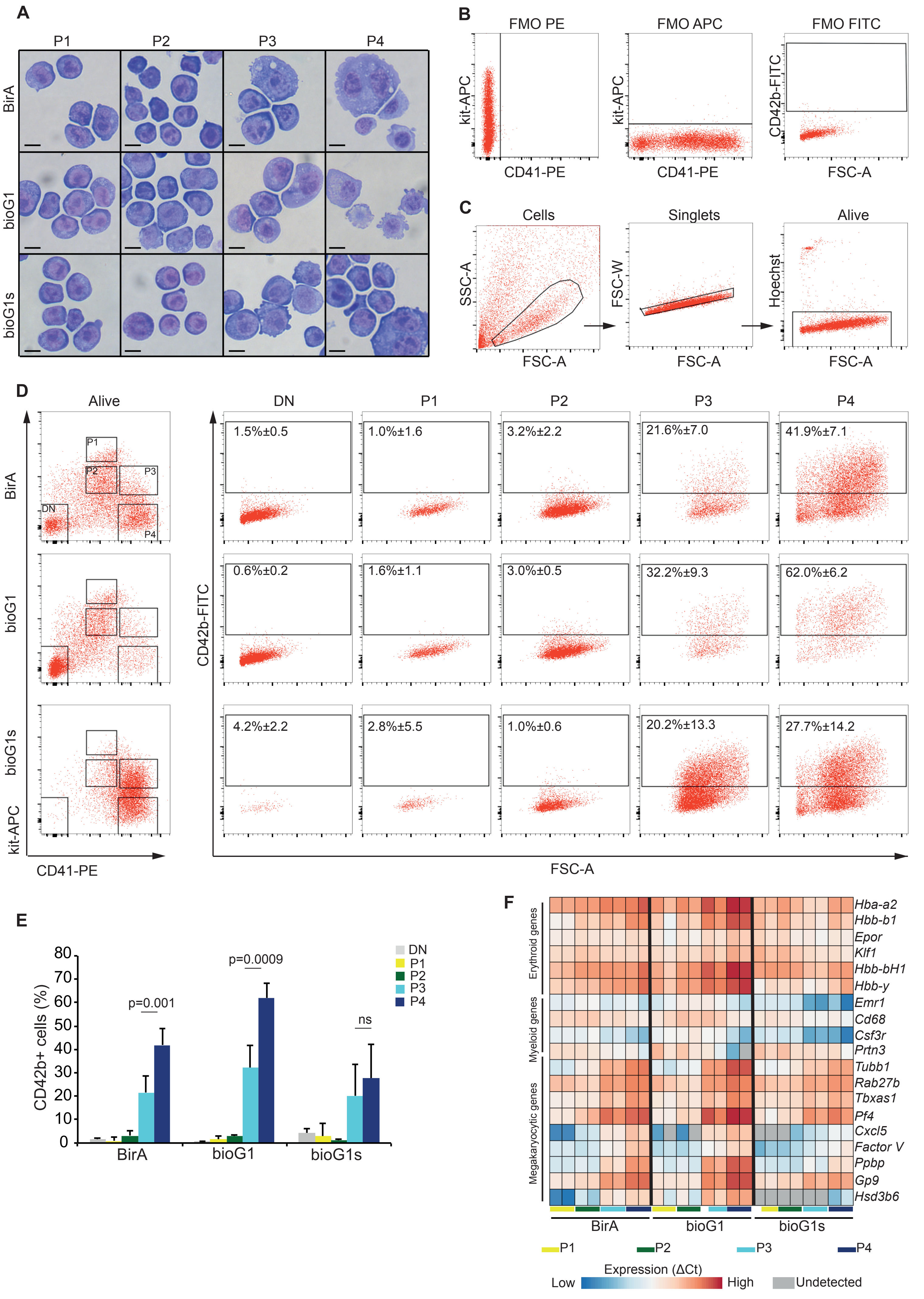

#### Supplemental Figure 6

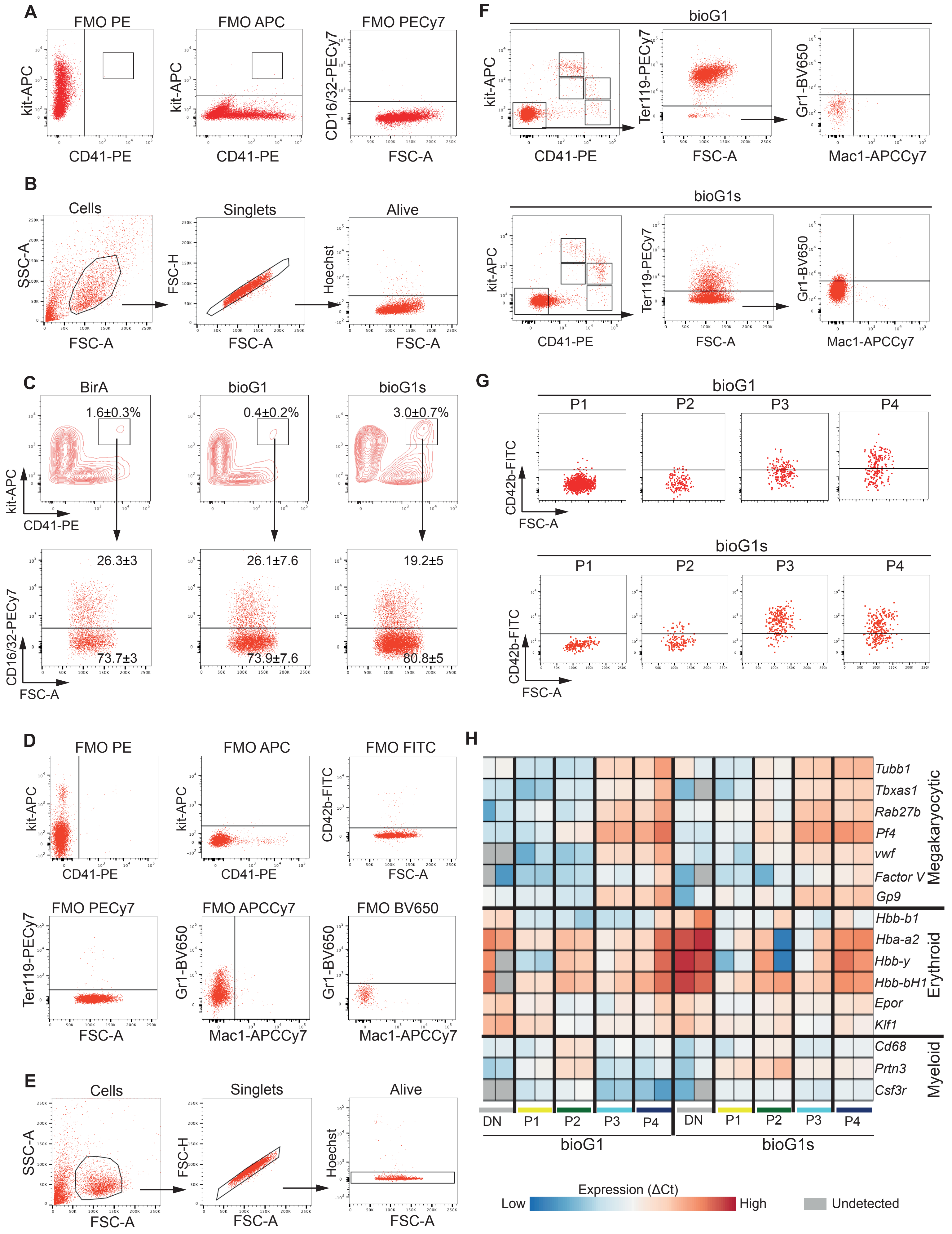

**Supplementary Table 1**

| Antigen | Conjugate | Supplier | Clone |
|---|---|---|---|
| CD11b (Mac1) | APCCy7 | BioLegend | M1/70 |
| CD16/32 | PECy7 | eBioscience | 93 |
| CD41 | PE | BioLegend | MWReg30 |
| CD41 | APC | BioLegend | MWReg30 |
| CD41 | BV421 | BioLegend | MWReg30 |
| CD42b | FITC | Emfret | Xia.G5 |
| CD61 | PE | BioLegend | 2C9.G2 (HMß3-1) |
| CD117 (kit) | PE | BioLegend | 2B8 |
| CD117 (kit) | APC | BioLegend | 2B8 |
| Gr1 | FITC | eBioscience | RB6-8C5 |
| Gr1 | BV650 | BioLegend | RB6-8C5 |
| Ter 119 | PECy7 | BD Pharmingen | TER-119 |
